# Topological spatial coding for rapid generalization in the hippocampal formation

**DOI:** 10.1101/2025.08.26.669250

**Authors:** Heejun Kim, Daniel C. McNamee, Nayeong Jeong, Sang Wan Lee

## Abstract

The brain navigates complex environments by combining entorhinal grid codes with hippocampal place codes. Although grid codes effectively represent a current environment’s geometry, their capacity to generalize across topologically analogous environments with different reward and state structures remains poorly understood. We introduce topology-aware grid coding (TAG), a computational theory that leverages topological invariance to generalize to new environments with the same topological structure but with different geometry. Drawing on the Euler characteristic from algebraic topology, TAG integrates complementary neural codes built on foundational grid bases: place codes serving as 0D vertices for self-localization, boundary codes acting as 1D edges to learn policy-independent grid codes through state prediction errors, and corner codes functioning as 2D faces for identifying topologically significant states. TAG grid codes remain stable under topology-preserving deformations yet discriminate among non-isomorphic structures. TAG develops policy-independent grid codes for novel structures more rapidly and robustly than existing approaches, balancing structure and policy encoding for multi-subgoal navigation without extensive planning. Finally, we show that TAG is compatible with transformer architectures, enabling its integration into scalable neural networks. Together, the TAG theory describes the essential nature of geometric objects to explain how the entorhinal-hippocampal system maps the unique topological structure of spaces.

## Introduction

Biological navigation and decision-making rely on representations of environmental structures. Specifically, grid cells exhibit periodic firing patterns that encode spatial relationships independent of specific locations^1^, while place cells uniquely identify distinct environmental states^2^. These neural representations jointly form a cognitive map, an internal representation of environmental structure that enables animals to predict, plan, and adapt to dynamic environments^3–6^. Interestingly, a subset of grid cells^7^ and place cells^8^ maintain firing patterns that are largely invariant to the animal’s moment-to-moment behavior policy, a feature that could allow the same neural map to support different tasks and reward configurations without retraining.

While these biological insights have inspired numerous reinforcement learning (RL) models, most rely on behavioral policy-dependent mechanisms for decision-making^9,10^. This requires the agent to relearn or transfer between policies when goals change. However, this approach undermines behavioral flexibility, since goal changes force the agent to switch to a different policy representation. This contrasts with biological systems that maintain a single, policy-independent cognitive map—making it biologically implausible.

Extending these neural observations, a growing body of work has used computational models to model policy-independent characteristics of cognitive maps. To overcome the shortcomings of policy-dependent learning, recent work has sought policy-independent map representations. Piray et al.^11,12^introduced the default representation (DR), offering a policy-regularized cognitive-map framework that supports flexible decision-making and accounts for the policy-independent firing patterns observed in grid and place cells, and demonstrated its compositional generalization by stitching together novel arrangements of familiar maze components. However, their applicability to more dynamic, multi-goal environments—closer to real-world biological navigation—remains unexplored, and their zero-shot generalization is confined to environments composed of identical object shapes.

Recent evidence suggests that these policy–independent spatial representations are not confined to encoding metric layouts but also capture topological features across hippocampal and connected brain regions, where they underpin flexible decision-making and generalization. Such topological encoding is observed in hippocampal ensembles^13,14^-including CA1 place cells^15,16^-which reliably reflect environmental topology across diverse geometries. Moreover, subicular neurons exhibit even stronger topology representations than those in CA1^16^. We suggest that these functional cell populations collectively encode cognitive maps in a *topologically invariant* format. Such abstract topological maps may guide choice-point decision-making^17^ and re-planning^18^ while remaining robust under topology-preserving transformations of an environment^19^. Potentially, such topological coding contributes to generalization across topologically isomorphic environments. Although these findings bolster the hypothesis that spatial structure is abstractly encoded in a topological manner, the precise neural coding mechanisms that give rise to these topological representations, and the computational mechanism on how they are leveraged for task generalization, remain poorly understood.

Building on place and grid codes, neurophysiological studies have identified boundary vector cells (BVCs) that fire at specific distances to walls^20^ and corner cells that respond at junctions of environmental boundaries^21,22^. As a computational effort to integrate these signals into the predictive map framework, BVC–SR uses BVC representations as basis functions for successor representation learning^23^. Moreover, subicular neurons have been shown to form generic spatial maps that generalize across distinct environments^24^. Despite these advances, how these subicular neural activities contribute to the formation of abstract topological state representations remains poorly understood.

In this work, we introduce topology-aware grid coding (TAG), a unified framework which provides a computational account of how the brain acquires topological representations, and leverages these representations for flexible decision-making and generalization. We hypothesize that behavior policy-independent grid coding is essential for topology-awareness. TAG integrates entorhinal grid codes, hippocampal place codes, and subicular boundary and corner codes into a single framework for learning the topology-aware grid representation. TAG operates through a double-loop architecture: the inner loop-driven by place and boundary codes-focuses on achieving policy-independent grid coding, while the outer loop-driven by boundary and corner codes-enhances topology-awareness. This design elucidates the role of topology-aware grid code as a basis for spanning the other neural codes and demonstrates how they collectively support both policy-independent grid representations and the extraction of environmental topology. We also show that TAG successfully predicts key neural findings related to the topological representations, underscoring its biological plausibility.

We demonstrate that the TAG grid code exhibits strong topology-awareness, remaining robust under topology-preserving environmental augmentations, such as shift, scale, and rotation of the environment. This robustness enables TAG to generalize across different environments sharing the same underlying topology. Moreover, TAG maintains sensitivity enough to distinguish between topologically non-isomorphic environments. Consequently, when exploring a topologically isomorphic environment, TAG naturally focuses on viable transitions—automatically avoiding any moves that would lead to non-existent, non-isomorphic connections—and thus achieves faster, more efficient coverage. This dual capability highlights the model’s flexibility in generalizing across isomorphic environments while accurately identifying meaningful topological differences. Leveraging its topology-awareness for decision-making, TAG achieves near-optimal performance in complex subgoal navigation tasks without the extensive planning or policy switching traditionally required in multi-subgoal reinforcement learning. Finally, we demonstrate that TAG can be expressed within a transformer architecture, positioning it as a potential building block for topology-aware deep learning models. This connection bridges biological principles with scalable AI systems, offering new pathways for both computational neuroscience and artificial intelligence research.

## Results

### Importance of topology-awareness in task generalization

In reinforcement learning (RL), an environment is typically modeled as a set of states and their transitions. However, it may be unrealistic for a biological agent to meticulously track all these transitions due to finite memory capacity. Storing every state transition while simultaneously adapting to new environments would impose a significant burden on memory. Additionally, the environment structure can be easily disrupted by minor changes in state transition dynamics, limiting the agent’s ability to generalize. This observation suggests that reducing sensitivity to minor changes could enhance generalizability, allowing the agent to focus on topologically significant features. We explore this idea using topological abstraction, in which states are classified into the topological state labels: junction, edge, and dead-end (Methods). This method simplifies any environmental structure into a representation that captures its essential topology, regardless of environmental scales and reward functions (Fig. 1a-c).

**Figure 1.**
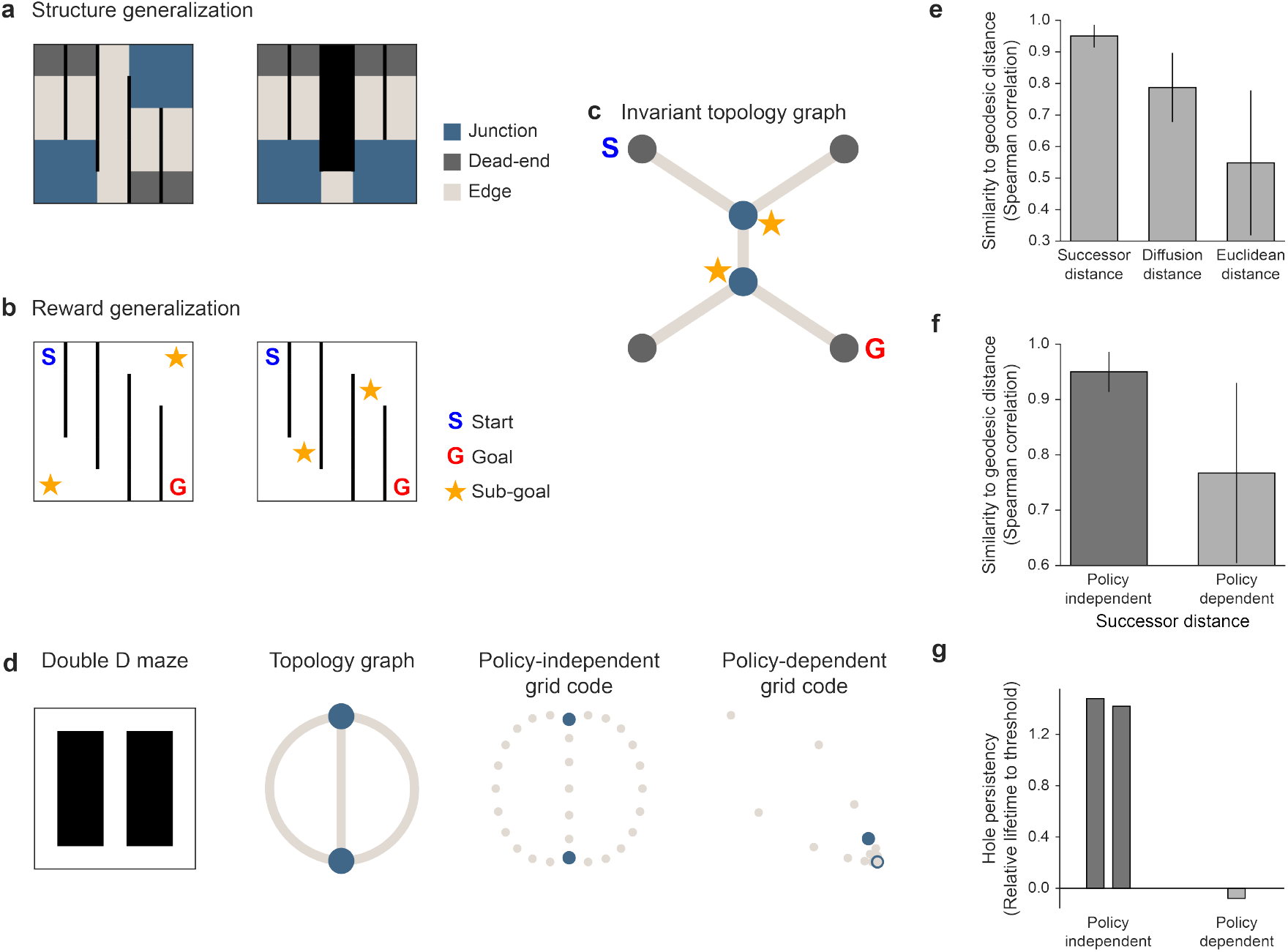
Topology-awareness of policy-independent grid code. **a**, Generalization across different topologically-similar structures. Three topological state labels—junction, dead-end, and edge—are used to abstract environment structures. **b**, Generalization across different reward locations associated with the same topological state label. **c**, Abstracted environment structures and reward locations can be represented as a graph, where nodes correspond to junction or dead-end state labels, and edges correspond to edge state labels. **d**, The policy-independent grid code aligns with the topology graph of the double D maze, demonstrating its topology-awareness, whereas the biased grid code does not. The grid code manifold visualizes the 2nd and 3rd axes of the grid code. **e**, Among various metrics, the successor distance best encodes the structure of the environment compared to the diffusion and Euclidean distances. Diffusion distance was calculated with the scale parameter *t* = 7, which empirically yielded the highest encoding score (refer to Supplementary Figure 3a). **f**, The environmental structure is effectively encoded only when the grid code is policy-independent with respect to the behavior policy. Error bars represent the standard deviation over 1000 randomly generated mazes. **g**, Persistent homology analysis^16^, a technique for detecting topological holes in manifolds, was applied to a double-D maze with two loops. To assess statistical significance, each hole’s lifetime (birth–death interval) was compared against a threshold defined by the 99.9th percentile of lifetimes obtained from randomly shuffled manifolds. In the resulting plot, this threshold is subtracted from each lifetime so that the y-axis zero point corresponds to the significance cutoff—intervals extending above zero thus reflect robust, statistically significant holes. The policy-independent grid code exhibits two prominent long-lived intervals rising above the threshold, consistent with the maze’s two-loop structure. In contrast, the biased grid code yields no intervals exceeding the threshold, suggesting a loss of true topological structure.

For structural generalization, the agent can adapt to distinct environment configurations with varying state transition dynamics, as long as they share the same underlying topology. This form of generalization abstracts environment structures by incorporating the connectivity patterns among the topological state labels, rather than the precise number or spatial arrangement of states. For instance, inferring the underlying topological structure shared between two state-spaces would facilitate cross-environment generalization (Fig. 1a, c). Similarly, reward generalization enables the agent to adapt to different reward placements within environments that belong to the same topological state label. In this context, the exact state associated with the reward becomes less crucial, provided the new location remains within the same topological state label. This abstraction helps streamline the decision-making by shifting the focus from re-identifying rewarding states to whether a rewarding state should be visited, and in what sequence the state levels should be traversed. As a result, the agent gains flexibility in adapting to changing reward locations (Fig. 1b, c).

### Policy-independent grid codes enhance topology awareness

Building on these observations, we explore the requirements for topology-aware structure representation. We focus on the successor representation (SR)^25^, a reinforcement learning model that predicts future state occupancy based on state-transition dynamics. The SR is known to capture hippocampal encoding of the structural and temporal dynamics of the environment.

Spectral decomposition of the SR matrix results in a grid code for encoding environmental structure^26,27^, from which one can infer relationships between states to learn diverse behavioral policies. To span a variety of policies, this basis necessarily encodes a policy-independent environmental structure. This aligns with the proto-value function^28,29^, which utilizes normalized graph Laplacian eigenvectors to represent the basis for state values under a similar philosophy.

The grid code of the policy-independent SR model corresponds to the graph Laplacian with equally weighted nodes (see Supplementary Note 1 for detailed derivation). In this case, the Laplacian eigenvectors embed nodes with similar connectivity closer together while separating those with weaker connections^30,31^. Such a characteristic ensures that states within the same topological state label (e.g., junctions or dead-ends) are closely embedded, whereas states across labels (e.g., distinct state labels connected by edges) are positioned farther apart. This implies that the SR grid code is suitable for representing the underlying topological structure of environments (Fig. 1d).

To evaluate the grid code’s topology embedding, we used the successor distance, a Euclidean distance on the successor coordinate constructed using SR eigenpairs (Methods). The successor distance was confirmed to represent environmental topology better than other distance metrics (Fig. 1e), including a simple Euclidean distance and the diffusion distance (Methods). It is ascribed to the successor distance’s ability to prioritize low spatial frequencies in spectral regularization^26^. We also confirmed that the successor distance can capture global structural patterns while being robust against small environmental perturbations (see Supplementary Note 5 and Supplementary Figure 2, 3b-c for more detailed analysis).

**Figure 2.**
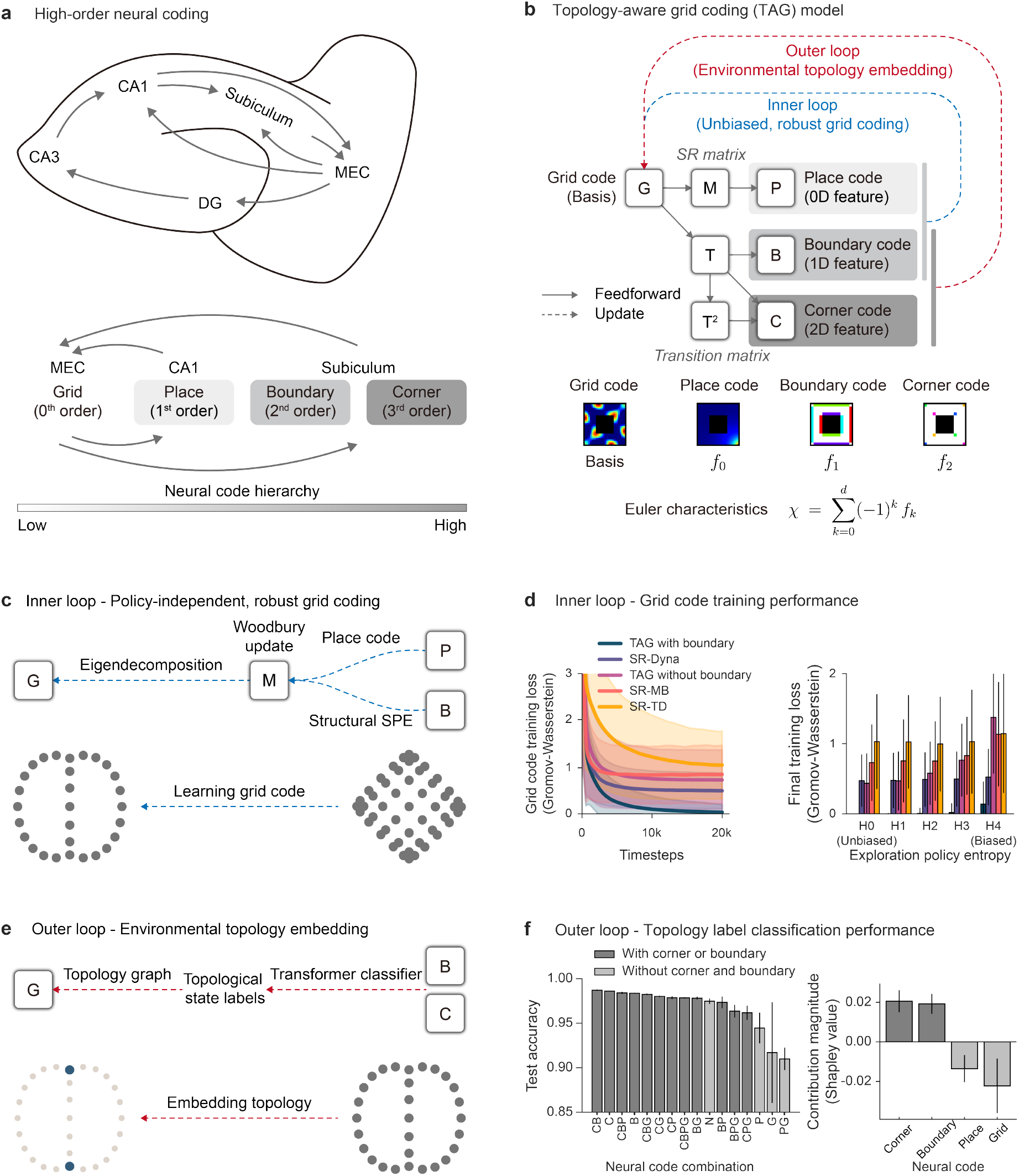
Topology-aware grid code (TAG) model. **a** (Top) Schematic of hippocampal circuitry depicting the sequential loop from the medial entorhinal cortex (MEC) through dentate gyrus (DG), CA3, CA1, and subiculum, and back to MEC. (Bottom) Hierarchical organization of grid, place, boundary, and corner codes, arranged in order along the neural processing stream. **b**, Inspired by the Euler characteristic, TAG model integrates grid, place, boundary, and corner codes into a single computational hierarchy, with each code contributing a distinct role in topology-awareness. These neural codes are updated using a double-loop update mechanism. **c**, Inner loop of TAG. The boundary code is utilized for computing the structural SPE, essential for learning policy-independent grid codes. **d**, (Left) Grid code training loss of models from rewardless random walk trajectories (Methods). (Right) Final training loss. Exploration policy entropy from H0 to H4 represents the policy bias from low to high (Methods). **e**, Outer loop of TAG. Boundary and corner codes are utilized as input to the transformer classifier to classify states into the topological state labels. **f**, (Left) Test accuracy of a transformer model for classifying the topological state label for each state given neural codes. “N” indicates that none of the neural codes are provided, with only blocks as input. (Right) Contributions of each neural code to topology classification (Methods). Shaded areas and error bars represent standard deviations across 1000 randomly generated mazes.

**Figure 3.**
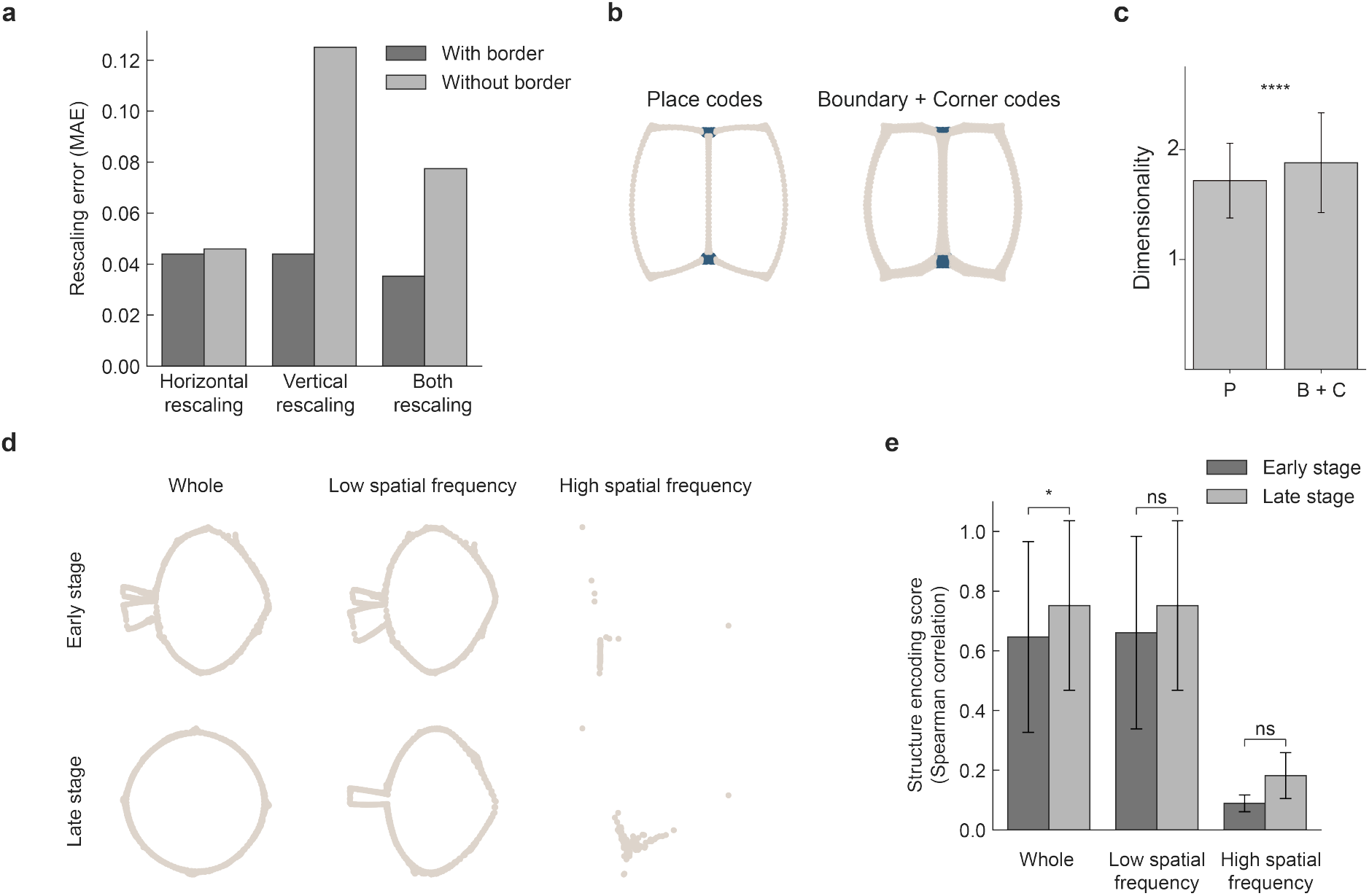
Neural predictions of TAG. **a**, Entorhinal grid cells rapidly adjust their spatial frequency to match environment size changes^44^. TAG with boundary code reproduces this rapid, accurate rescaling. TAG without boundary code shows biased grid scaling and high error rates. **b**, Isomap of TAG place and boundary–corner codes reproduces the CA1 and subicular topology embedding. **c**, TAG boundary–corner manifold shows greater dimensionality than TAG place code manifold. Error bars represent the standard deviations of 1,000 randomly generated environments. **d**, CA1 manifold topology similarity increases significantly across training stages, with strongly spatial (SS) and weakly spatial (WS) subpopulations showing little change^15^. TAG place code manifold topology similarity replicates this training stage-dependent increase, while SS/WS submanifolds—derived by thresholding SR eigenvalues into low- and high-frequency bands—remain largely unchanged. **e**, CA1 population manifold shows a significant stage-to-stage increase in topology similarity. TAG place code manifold similarly exhibits a significant increase, whereas high and low spatial frequency submanifolds do not. Error bars indicate the standard error across seeds.

While the grid codes are shown to be effective in topology embedding, it can be disrupted by a biased policy, such as the one associated with a reward-maximizing value function. The policy bias alters the weight distribution of the adjacency matrix, even though the row vector sums remain unchanged. This distributional shift affects the eigenvectors, impairing their ability to accurately encode structural relationships. This occurs because it perturbs the successor coordinates and distorts the manifold structure. As a result, the low-dimensional embedding of the SR grid code deviates from the true topology, culminating in structural encoding that is misaligned with the geodesic distance (Fig. 1d, f, g). For instance, biased policies could prioritize specific branches, distorting the actual distances between states and compromising the fidelity of structural encoding.

These observations suggest that by leveraging policy-independent successor distances, we can construct biologically plausible topology embeddings, for example, with the edge weights of the topology graph determined by their successor distances. This approach integrates the spatial relationships encoded in the SR grid code into a graph representation, enabling a natural and biologically inspired framework for topology embedding.

### Topology-aware grid coding theory (TAG)

Anatomical and tracing studies have revealed multiple interlocking recurrent circuits linking the medial entorhinal cortex (MEC) with hippocampal output regions^32–35^. First, direct reciprocal projections connect MEC with CA1 and MEC with subiculum (SUB), giving rise to two distinct loops—a MEC–CA1–MEC loop and a MEC–SUB–MEC loop—each capable of supporting recurrent processing. Second, the classic trisynaptic circuit routes from MEC through dentate gyrus (DG), CA3, and CA1 to SUB before looping back to MEC, establishing a processing gradient across regions, with activity progressively transforming as it travels from MEC to CA1, and finally to SUB. This organization suggests that grid cell inputs in MEC give rise to place cell representations in CA1, which are further elaborated into boundary-and corner-selective representations in subiculum^20,21^, corresponding to successive orders of spatial structure (Fig. 2a). By aligning each stage of our four-layer model with these well-characterized pathways, we ground grid, place, boundary, and corner codes in their native anatomical substrates and motivate their hierarchical organization.

Based on these observations, we hypothesize that the grid code serves as a foundation for constructing a topological hierarchy, encompassing hippocampal place cells, subicular boundary cells^20^, and subicular corner cells^21^. To systematically investigate this hypothesis, we developed a topology-aware grid coding (TAG) model (Fig. 2b) with four distinct layers: a grid, place, boundary, and corner code. Our architecture introduces three key innovations. First, each code contributes uniquely to topology awareness: the grid code constructs an environmental structure, the place code enables self-localization, and the boundary and the corner code encode the first- and second-order environmental constraints. Second, all codes are derived from the grid code basis, creating a unified computational hierarchy (see the computational graph in Fig. 2b). Third, this layered architecture aligns with the mathematical principles of Euler characteristic^36^, which measure topological invariants by enumerating 0-, 1-, and 2-dimensional features and beyond. It not only suggests potential topological invariance but also predicts higher-order codes. For example, the place, boundary, and corner codes correspond to 0-, 1-, and 2-faces in geometry. Extending further, deep-layered codes could implement *k* -faces.

The grid code of TAG decodes both policy and structural representations, reflecting the activities of hippocampal place cells and subicular boundary and corner cells. This is accomplished through computation of both the SR matrix and the state-transition matrix, where each matrix’s eigenvalues are multiplied with the grid code (See Supplementary Note 1 for details). This framework models the neural pathways connecting the medial entorhinal cortex (MEC) to the hippocampus (HP), as well as the connection from the MEC to the subiculum. Next, the place code of TAG is modeled as the column vectors of the SR matrix, as extensively discussed in prior studies^11,12,26,37^. This modeling approach highlights the role of the place code in optimal goal-directed planning^11^.

The model incorporates boundary codes by leveraging the state-transition matrix. This setting allows identifying environmental boundary constraints where single-step transitions to adjacent states are impossible. Moreover, because boundary cell activation is direction sensitive^20^, our model explicitly encodes boundary directionality.

The model also incorporates the corner code, which is activated near the intersection of two borders^21,22^, by combining the boundary code with the squared transition matrix that represents the probabilities of two-step transitions. This approach effectively identifies both concave and convex corners that constrain the state transition. In particular, the corner code activates when two-step transitions are impossible, while one-step transitions to adjacent directions are both possible or both impossible. This transition constraint aligns with the experimental findings that corner cells remain inactive when borders are disconnected^21^, as such configurations permit two-step transitions.

Note that the corner code, like the boundary code, incorporates directional information to enhance spatial encoding precision. While direct evidence for allocentric direction sensitivity in corner cells remains to be demonstrated, our model draws inspiration from the retrosplenial cortex’s egocentric vertex cells^22^, which exhibit head direction-dependent firing patterns. In line with this finding, our model exhibits direction-selective corner cell activation patterns, potentially enabling the integration of egocentric and allocentric representations for more accurate spatial encoding.

The TAG model learns its own topological basis, the grid code, through dynamic interactions with the environment. This learning process unfolds across two distinct levels: an inner loop that distills state-transition experiences into a policy-independent basis (Fig. 2b, c) and an outer loop that distills environmental variations into a task-independent basis (Fig. 2b, e). While the inner loop enhances the grid code’s robustness against policy bias within individual tasks, the outer loop expands the grid code basis across multiple tasks. These dual update mechanisms, operating at different timescales, mirror the fast-slow learning principles in neuroscience and AI^38–40^.

### TAG inner loop - policy-independent, robust grid coding

The inner loop of TAG uses the place and boundary code for robust learning of state-transition dynamics (Fig. 2c). While model-based reinforcement learning approaches, including TAG, learn state-transition dynamics through state prediction error (SPE) - which measures discrepancies in transition probabilities - optimal learning requires policy-independent updates that capture transition probability differences under random-walk policies. We propose that the boundary code can be used to compute an SPE under the unbiased policy, which we term the *structural SPE*.

The structural SPE is defined as the difference between the random-walk probability distribution before and after exploration steps, where its probability distribution across adjacent states *s* is estimated by applying a softmax function to the inverted boundary code *B* (where 0s and 1s are flipped) (structural SPE; Fig. 2c):

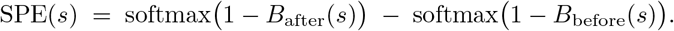

The agent then uses this information to update the transition matrix and the SR. While maintaining and inverting the full transition matrix would be both computationally intensive and biologically implausible, the Woodbury update method offers an efficient alternative by enabling rank-one updates without requiring full matrix inversion^11^. Using this method, the agent integrates place code and structural SPE for efficient updates, followed by eigendecomposition to calculate the grid code, thereby enabling robust encoding of the updated environment structure. Incorporating hippocampal place code to update the entorhinal grid code reflects the hippocampus-to-entorhinal cortex feedback pathway^41^. Refer to Supplementary Note 2 for full algorithmic details.

To assess the efficiency of the boundary code-based learning rule, we evaluated a TAG agent across 1,000 randomly generated environments using rewardless trajectories sampled through random walks over the entire action space (Methods). We tested the model’s resilience to exploration policy bias by systematically varying bias levels during testing. The TAG model incorporating the boundary code demonstrated superior learning efficiency and accuracy (Fig. 2d). This is attributable to two key mechanisms: the SPE signal enables the one-shot learning capability by facilitating efficient SR updates, and the boundary code corrects for exploration policy bias through structural SPE. Notably, while increased policy bias slightly reduced learning speed, the TAG model shows reliable final performance even under highly biased exploration conditions.

We compared the TAG model’s performance with several alternative approaches. SR-Dyna^37^ attempts to mitigate policy bias to some extent through random-walk replay, but this correction does not significantly enhance grid code learning. The TAG model without boundary code, while learning more rapidly, inherits exploration policy bias and exhibits higher training loss than SR-Dyna, underscoring the boundary code’s crucial role in bias mitigation. SR-MB^37^ achieves rapid initial learning through direct transition matrix updates using the Woodbury matrix identity^11^, but like TAG without boundary code, it remains susceptible to policy bias, resulting in biased convergence. This reinforces the necessity of policy-bias correction for accurate grid code learning. SR-TD^25,37,42^, which relies exclusively on iterative updates, demonstrated the slowest learning performance. Without mechanisms like random-walk replay, it struggled to adapt and remained highly vulnerable to policy bias.

Taken together, only the TAG model with boundary code-based learning achieved a low training loss under biased exploration policies. This robustness highlights the essential role of the boundary code in ensuring accurate and efficient learning, even in the presence of exploration policy bias.

### TAG outer loop - environmental topology embedding

Our TAG theory predicts that while the lower layers of the TAG serve to learn a policy-independent environmental topology (Fig. 2e), the higher layer of the TAG hierarchy, the subicular boundary and corner code, affords the agent the leverage to learn a task-independent environmental topology. To test this idea, the outer loop of TAG uses the boundary and corner codes to refine the grid code basis over multiple tasks (Fig. 2c). Specifically, we examined the ability of the corner code to classify states into distinct topological state labels (junctions, dead-ends, and edges).

A transformer-based architecture^43^ was used for topology classification (Fig. 2e). The model takes as input combinations of various neural codes (corner, boundary, place, and grid) along with block information, and then categorizes them into three topology labels (junction, dead-end, or edge) at the feedforward layer (Methods). To evaluate the contribution of each neural code, we systematically assessed the classifier’s test accuracy across all possible input feature combinations.

We found that higher layers contributed progressively more to topology embedding, with the boundary and corner code demonstrating pronounced effects. Ablation studies, in which the boundary or corner code is removed from the model, confirmed their contribution to topology classification (Fig. 2f). This influence likely stems from the boundary and corner code’s direction-sensitive nature to indicate the presence or absence of obstacles in distinct directions. In contrast, the TAG’s lower layers, the place and grid codes, showed negative contributions. Their performance degradation is because the place codes are often task-dependent and goal-based, while the grid codes are sign-variant, exhibiting inconsistent encoding of positive and negative regions. This pattern of contribution reflects the hierarchical organization of the neural codes.

Building on our earlier discussion, successor distance-based topology graphs provide a means to determine the edge weights between nodes. By leveraging our classification results, we identified distinct topological state labels-junctions, dead-ends, and edges-which serve as nodes and edges in the topology graphs. This approach enables the construction of a biologically plausible topological embedding of the environment, aligning with its structural constraints observed in neural representations.

### Neural predictions of TAG

The TAG framework yields a series of testable predictions about hippocampal–entorhinal activity. First, we examined whether TAG can account for the adaptive rescaling of grid cell firing fields following the environmental size changes^44^. Agents performed unbiased random-walk exploration within the resized open field to path-integrate its new geometry (Methods). TAG equipped with the boundary code accurately calibrated its grid scale to match the resized environment—reproducing the neural rescaling effect (Fig. 3a). In contrast, TAG lacking the boundary code failed to replicate this effect. Although a randomwalk policy can generate a policy-independent map, it proved insufficient for capturing the precise scaling profile on its own, highlighting the indispensable role of the boundary code. These results demonstrate that structural SPE, computed via the boundary code, supports rapid and policy-independent adaptation of the grid code to environmental geometry changes.

Second, we evaluated whether TAG’s place codes and combined boundary–corner codes recover the topological manifolds observed in CA1 and subicular neuron populations, and whether they account for the characteristic dimensionality differences between these regions. Specifically, we sought to replicate recent findings showing that CA1 population activity encodes environmental topology, with isomap projections of CA1 firing patterns faithfully reflecting the geometric structure of the square-donut arena^15,16^. TAG place codes successfully reproduce these topological embeddings in low-dimensional projections across the same variety of environments (Fig. 3b), highlighting the generality and robustness of TAG’s topology representation.

Beyond CA1, recent studies have shown that subicular neuron populations also encode environmental topology via low-dimensional embeddings^16^. In TAG, the combined boundary and corner codes are designed to emulate these subicular representations. To model this, we concatenated the TAG boundary and corner vectors, applied a geodesic distance–dependent decay to their activations, and then conducted dimensionality reduction (Methods). This distance-dependent attenuation—reflecting the gradual decay observed in biological boundary vector cells^20^ and corner cells^21^. The resulting embeddings closely reproduced the environment’s topological layout (Fig. 3b), demonstrating that TAG’s boundary–corner population captures the topology-encoding dynamics of the subiculum.

Another interesting characteristic of the subiculum is that its populations exhibit significantly higher manifold dimensionality than CA1^16^. This pattern is successfully reproduced by our TAG model (Fig. 3c). We attribute this effect to the enriched geometric information encoded by boundary and corner codes, which enhance the topology embeddings beyond the capacity of place codes alone. To explore how key parameters shape this effect, we performed a parameter sweep over the SR discount factor *γ*_*SR*_ and a boundary-corner decay rate *γ*_*BC*_, which governs the geodesic attenuation of boundary + corner activations. Topology encoding was quantified using the similarity to geodesic distance across these parameter combinations (Methods), revealing optimal structure encoding at *γ*_*SR*_ = *γ*_*BC*_ = 0.99. These findings highlight that robust topological representations depend not only on place codes within CA1, but also critically on subicular contributions—boundary and corner codes—to fully capture environmental geometry.

Finally, we examined TAG’s account of how strongly and weakly spatial subpopulations jointly contribute to the formation of topology representations across learning stages. Manifold analyses of hippocampal CA1 have revealed that strongly spatial (SS) and weakly spatial (WS) subpopulations contribute differently to topology formation^15^. In TAG, SS place codes correspond to SR constructed with eigenvectors with large eigenvalues, whereas WS place codes correspond to those constructed with small eigenvalued eigenvectors—reflecting that eigenvectors with large eigenvalues capture global structure with low spatial frequencies, while those with small eigenvalues encode local features with high spatial frequencies (see Supplementary Note 5). Early and later training stages were designed to reflect earlier and more advanced stages of learning (Methods). Guo et al. reported that SS manifolds already exhibit robust topological embeddings across training stage, WS manifolds alone convey little topology, but the combined SS+WS manifold shows a statistically significant training stage-dependent increase in topological fidelity. The TAG model successfully recapitulates these observations: both SS and WS submanifolds exhibit training stage-specific effects, while the combined boundary–corner manifold yields the most complete topological embedding (Fig. 3d-e). Notably, while a spectral regularization regime often recommends discarding low-eigenvalue, high spatial frequency components as noise^26^, our findings demonstrate that WS components—despite their higher spatial frequency—are essential for constructing accurate topological representations. Indeed, in the later training stage the manifold reconstructed with WS components (Fig. 3k) reveals a far more complete topological structure than models relying solely on high-eigenvalue modes.

### Robustness of TAG grid code across topology-preserving environment augmentations

To achieve rapid generalization across tasks, agents should first understand the environment’s fundamental topological patterns (such as junctions and dead-ends) before learning precise state-transition dynamics. To evaluate the TAG grid code’s ability to capture these underlying topological structures, we introduced environmental augmentations that modified state-transition dynamics while maintaining the underlying topology. We then measured TAG’s performance across various topology-preserving environment augmentations: noise, shift, scale, and rotation (Methods).

We evaluated whether the TAG grid code captures and preserves the underlying topology across augmented environments (Methods). Notably, the grid codes obtained from differently augmented environments aligned almost perfectly. This result suggests that the low-dimensional representation of the grid code effectively preserves the topology, even when the high-dimensional manifold undergoes transformations such as rotation or scaling (Fig. 4b).

**Figure 4.**
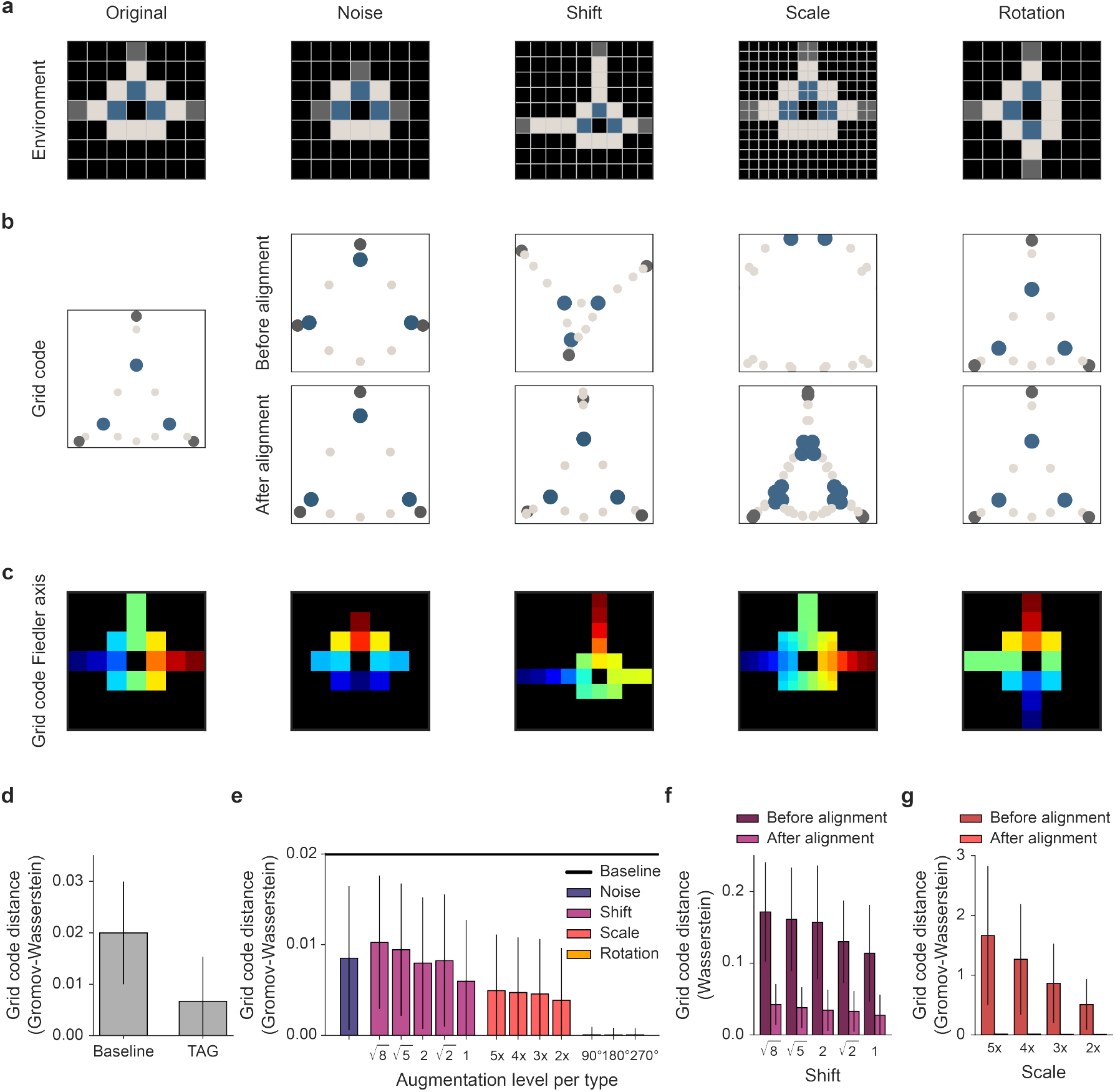
Robustness of TAG grid code across topology-preserving environment augmentations. **a**, Illustrations of topology-preserving environment augmentations. Noise augmentation involves randomly adding blocks while preserving the underlying topology. The shift augmentation shifts the environment, with the illustrated example showing a shift of factors *x* = 2 and *y* = 2. The scale augmentation enlarges the environment, demonstrated with a scaling factor of *k* = 2. Rotation augmentation rotates the environment, with the example corresponding to a rotation factor *t* = 1. **b**, The visualization of the top two axes of the successor coordinate before and after alignment using the Procrustes algorithm. Block states are omitted to prevent zero-value alignments that hinder precision. **c**, Visualization showing the first axis of the successor coordinate, with red representing positive values and blue representing negative values. **d**, Robustness of TAG grid code represented as a GW distance between the original and augmented grid codes, compared with a baseline representing GW distances between grid codes from non-isomorphic environment pairs (Methods). Analysis is conducted using the top 10 dimensions of the successor coordinate. **e** Robustness of TAG grid code demonstrated in detail for each augmentation type and level. Augmentation levels are quantified based on specific metrics: Euclidean distance for the shift augmentation, scaled amount for the scale augmentation, and rotation angle for the rotation augmentation. **f-g**, Grid code distances before and after alignment for shift (f) and scale (g) alignments. In (f), Wasserstein distance is used as it is rotation-variant, making it suitable for assessing the effect of rotation alignment. Error bars represent the standard deviations obtained from 1000 randomly generated mazes.

On the other hand, conventional structural representations relying on single-axis information, such as ones using the second-largest eigenvector of the graph Laplacian or SR for subgoal or option discovery—like eigenoptions^45^, normalized cuts^26,46^, or bottleneck encoding methods^47^—struggle to capture the inherent topology and are easily perturbed, especially under noise or shift augmentations (Fig. 4c). Furthermore, in scale augmentations, where the grid code manifold expands, this limitation becomes even more pronounced. Stachenfeld et al.^26^ also highlighted that Fourier interpolation is not suitable for grid code representations, as it tends to blur critical boundaries. Similarly, in this context, Fourier interpolation fails to preserve the environmental boundaries, underscoring the need for alternative regularization strategies. Incorporating boundary codes during interpolation can address this issue by maintaining boundary information while interpolating other aspects of the representation.

To further confirm our observations, we measured the distances between the original TAG grid code and that of augmented environments. First, across all augmentations, the distances remained consistently low compared to the baseline distance measured between grid codes of non-isomorphic environments (Fig. 4d), confirming the robustness of the TAG grid codes under various augmentations. Second, detailed analyses for each type of environmental augmentation revealed the following findings (Fig. 4e): for noise augmentation, the grid code distances confirmed that topology-preserving random perturbations do not significantly alter the grid code’s topological structure. For shift augmentation, certain geodesic distances between topological state labels are increased with the shifting distance, reflected in the grid code distances. Despite these changes, the overall topology remained unaffected. Shift augmentation caused a rotation of the entire manifold in space, visually evident in Fig. 4b, while explaining the perturbations in Fig. 4c. This is quantitatively confirmed by the grid code distance being significantly reduced after the alignment (Fig. 4f). For scale augmentation, while geodesic distances of edges increased with scaling, the topological structure remained intact. Grid code distances were significantly reduced after size alignment (Fig. 4g), underscoring the need for the alignment when evaluating topological similarity. This adaptation is analogous to the rescaling of grid codes^44^, potentially acting as a neural mechanism that allows rapid adaptation to isomorphic environments with different scales. Finally, grid codes were completely invariant to rotation augmentation, as it merely re-indexed states without compromising underlying structural representations.

### Topological structural generalization

The grid code’s topological invariance (Fig. 4) suggests the TAG model’s potential to adapt and generalize across topologically isomorphic environments. Although isomorphic environments share the same topological structures, their diverse state configurations and layouts pose significant obstacles to effective generalization. To address this challenge, TAG constructs a topology-abstracted representation of the environment, which it then uses to “decode” navigation policies by mapping this shared topological graph onto a novel configuration (Fig. 5a–b). This approach enables efficient zero-shot transfer between isomorphic environments.

**Figure 5.**
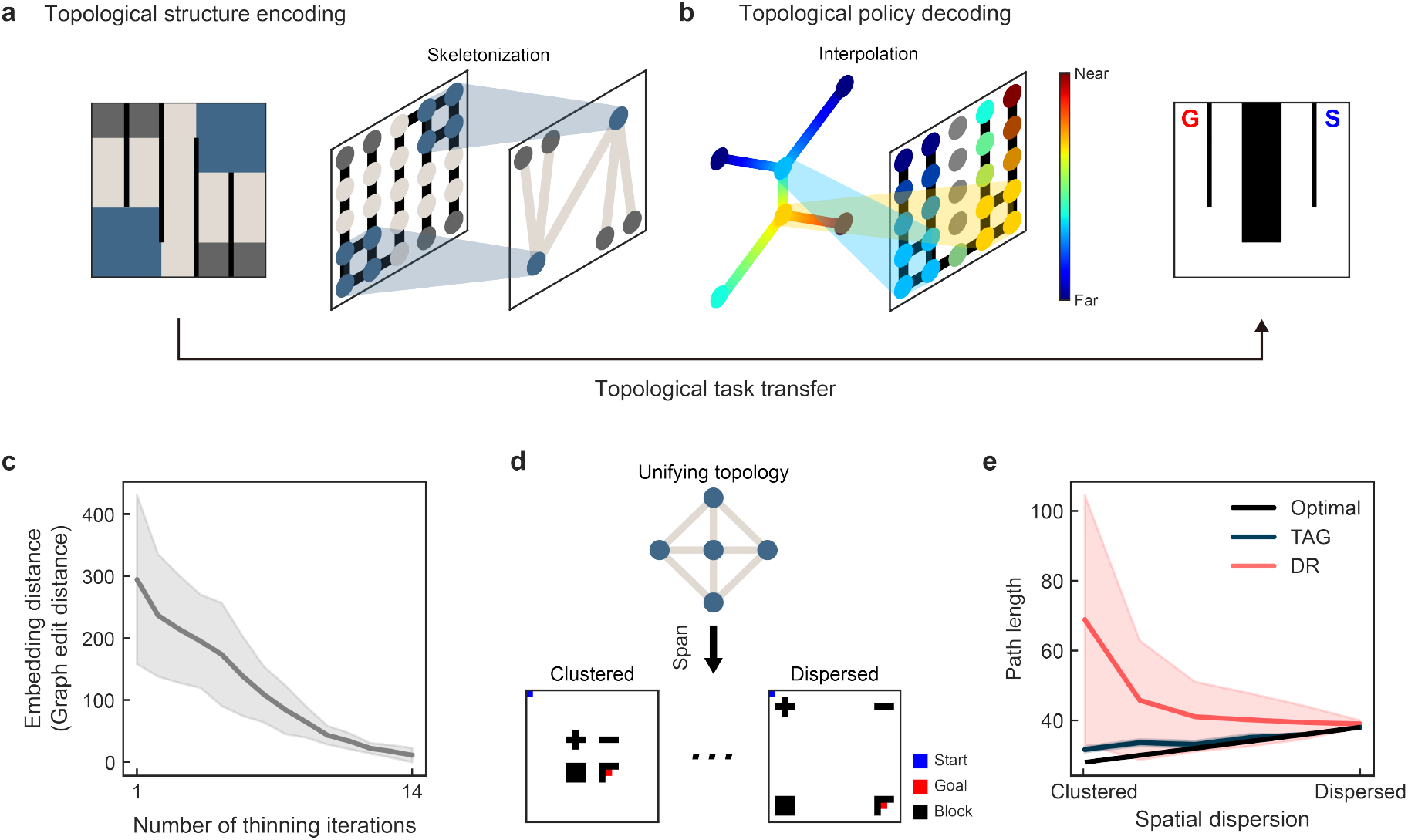
Topological structure generalization. **a, c**, Topological structure encoding pipeline: **a**, The agent embeds the environment structure into a topological abstract graph by skeletonizing the environment using a thinning algorithm (Methods). **c** Embedding distance between skeleton graphs of two isomorphic mazes decreases with more thinning iterations (Methods). Because environments differ in the total number of thinning iterations required for skeletonization, each trajectory was linearly interpolated to a uniform length of 14 steps (the maximum observed) to enable direct comparison. **b, d, e** Topological policy decoding pipeline: **b** In a novel isomorphic maze, the agent computes the shortest distance on the topology graph and projects it back onto the layout via kernel interpolation. **d** Policies for environments ranging from clustered to dispersed block configurations can all be decoded from a single unifying topology, as these environments share the same underlying topological structure. **e** TAG’s topology-guided decoding outperforms the DR approach^12^, which builds predictive maps by summing individual block influences.

Topological encoding refers to the abstraction of an environment’s structural layout into its underlying topological graph. As shown in Figs. 1 and 2, this process begins with skeletonization of the environment using a thinning algorithm (Methods). Each non-block state is then represented as a graph node, categorized as either a junction or dead-end, with connections between them forming graph edges (Fig. 5a). Biologically, this abstraction parallels subicular coding, which leverages boundary and corner codes to detect environmental features such as walls and corners—a process implemented in TAG’s outer loop for topology identification (Fig. 2b,e). Importantly, as thinning iterations progress, embeddings of isomorphic environments converge, demonstrating that the topological representation remains invariant across structurally equivalent mazes (Fig. 5c).

Building on our topological abstraction, we propose that shortest-path distances computed on the skeleton graph capture invariant structural relationships shared across all isomorphic environments, and that these distances can be zero-shot interpolated onto a novel layout to estimate optimal paths without additional learning. As illustrated in Fig. 5b, we first extract the maze’s skeleton graph, distilling the environment down to its junction and dead-end topology. Next, we compute a heat kernel over the graph nodes to quantify pairwise structural similarities. We then perform a three-stage composition to transfer shortest-path profiles back to the environment states: a radial basis function (RBF)-based projection from states onto skeleton nodes, propagation of distances among skeleton nodes via the heat kernel, and a second RBF interpolation from skeleton nodes back to states. This kernel-based interpolation of graph-level distance profiles enables zero-shot shortest-path estimation in previously unseen, topologically equivalent environments.

Although our topological policy decoding method naturally extends to any topologically isomorphic maze, it is instructive to compare it to the default representation (DR). Piray et al.^11^ originally introduced a Woodbury-based update rule for DR that adds one obstacle at a time, but because each update depends on the previous map, it fails to update in a truly compositional fashion. To address this, Piray et al.^12^ developed the predictive object representation (POR) framework, which compositionally infers a combined predictive map over multiple objects. However, POR’s zero-shot inference error grows sharply as obstacles cluster more densely, undermining accurate navigation without retraining. This compositional generalization applies to mazes built by shifting and rotating obstacle “objects”—ranging from tightly clustered to widely dispersed layouts—all of which share the same underlying topology (Fig. 5d). In contrast, our TAG framework directly decodes this shared topological structure to infer optimal paths in novel layouts without any additional learning.

To compare DR’s zero-shot compositional generalization with TAG’s topology-driven policy decoding, we evaluated navigation path lengths across a range of obstacle configurations (Fig. 5e). DR often approximates optimal path lengths in dispersed settings; however, as obstacles become more clustered, its compositional predictive map suffers from greater approximation error, leading to increased deviations from optimality. These inaccurate maps also result in high variability during stochastic navigation trials. By contrast, TAG consistently yields accurate path lengths across both clustered and dispersed layouts, exhibiting low variance across repeated stochastic trials.

Since POR focuses on object identities and their local dynamics, it cannot be extended to general topologically isomorphic environments. TAG, on the other hand, can minimize predictive errors while generalizing robustly to any topologically isomorphic configuration—including those beyond the compositional regime—by abstracting environmental connectivity rather than relying on object-level composition.

### Topology-awareness enables rapid exploration

Although knowing the exact environmental structure enables structural generalization to new layouts, the core challenge emerges from the complexity of mapping structural representations between environments—topological isomorphism alone does not guarantee straightforward state-to-state correspondence. Instead, agents must engage in active exploration and learning through direct environmental interaction. We investigated this hypothesis by evaluating an agent’s exploration and adaptation performance when provided with topological structure as prior knowledge, using exploration efficiency as our key metric for evaluating how well agents navigate the complexities of state correspondence (Fig. 6a). These insights led to a topology-aware exploration strategy for our TAG model (Methods).

**Figure 6.**
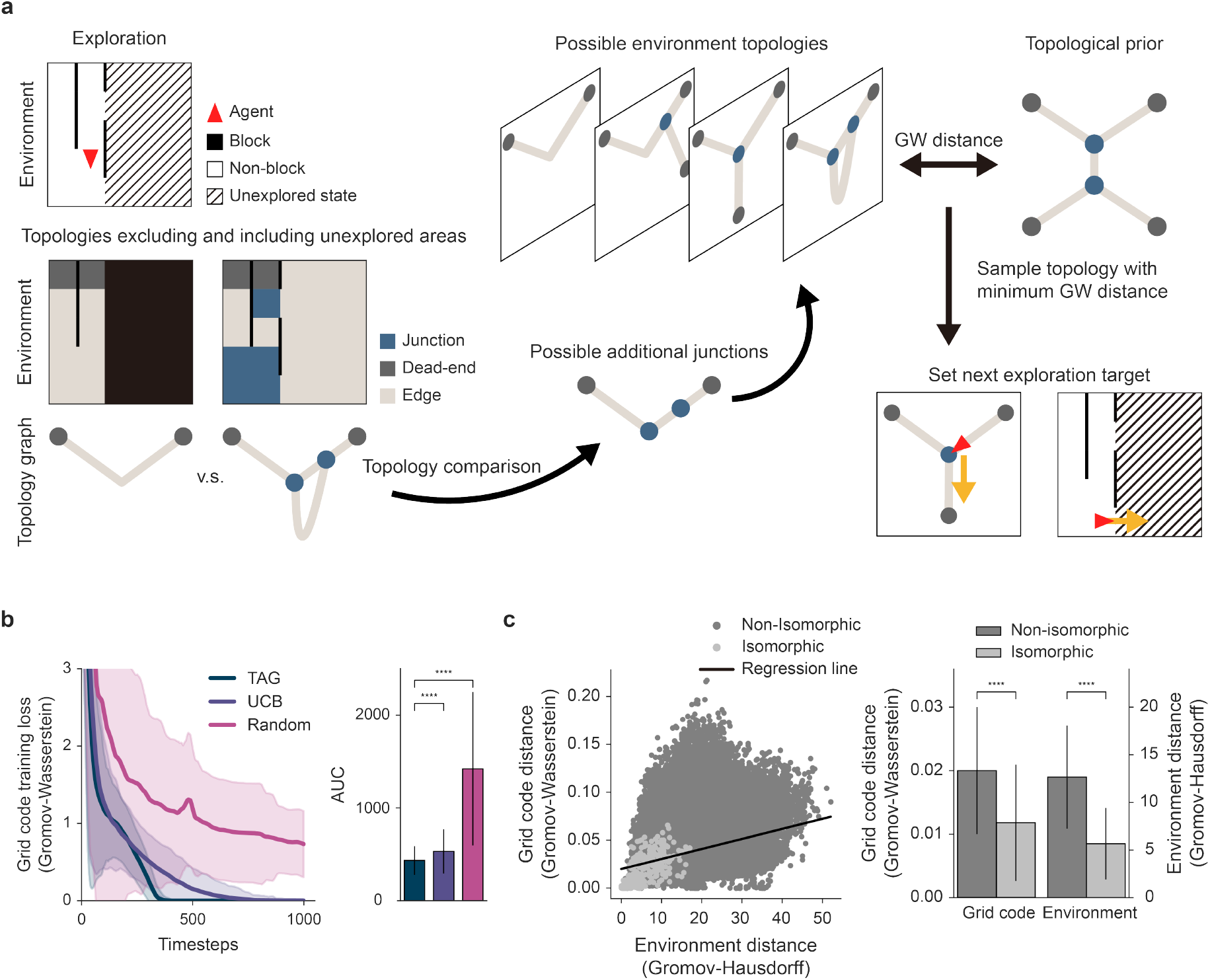
Topological exploration. **a**, The exploration algorithm identifies new topological edges by comparing topology graphs that exclude and include unexplored areas to find potential additional junctions. It generates candidate topology graphs by considering all combinations of possible new junctions and selects the graph with the minimum GW distance to the given topological prior. The next exploration target is set as the closest new edge from the agent’s location. **b**, Performance comparison of the proposed exploration algorithm against baselines, summarized using the area under the curve (AUC). **c**, Gromov-Wasserstein (GW) distances for grid codes and Gromov-Hausdorff (GH) distances for environment structures are evaluated across all distinct pairs of 1000 randomly generated environments. Each dot represents a pair, illustrating the relationship between grid code similarity (GW distance) and environmental structural similarity (GH distance) (Methods). The regression line shows a positive correlation between these two similarity metrics (*R*^2^ = 0.1). Shaded areas and error bars represent the standard deviations across 1000 mazes. ^****^ denotes statistical significance at *p <* 0.0001.

We evaluated TAG’s topology-aware exploration against two baseline approaches: a UCB (Upper Confidence Bound)-based exploration that prioritizes the least-visited states among the agent’s immediate neighbors, and random exploration. Our approach outperforms both baselines, highlighting the advantages of incorporating topological priors in guided exploration (Fig. 6b). The exploration process unfolds in two distinct phases. Initially, when no specific topological target has been identified, our method defaults to the UCB-based strategy, selecting the least-visited neighboring states. This early-phase behavior results in grid code loss comparable to the UCB-based baseline. However, once topological exploration targets emerge, the method demonstrates marked improvement in efficiency by transitioning from UCB-based exploration to strategic identification and pursuit of new topological edges based on partial topology information. This transition enables the agent to prioritize regions likely to contain novel topological features, significantly accelerating the exploration process compared to baseline approaches.

To further validate topology awareness of TAG, we empirically test whether distances between grid codes and distances between environments consistently distinguish isomorphic from non-isomorphic environment pairs (Fig. 6c). Results suggest that grid code distances successfully separate isomorphic and non-isomorphic pairs, mirroring the distinctions captured by environment distances. The state-wise structural distance between environments was less topology-aware than the TAG grid code distance, reinforcing our focus on environmental topology (See Supplementary Note 6 and Supplementary Figure 4 for details). Importantly, while the relationship between grid code distances and environment distances is not perfectly linear, their patterns align closely. This alignment demonstrates that the TAG grid code’s encoding of topology mirrors the structural relationships of the environment. Rare instances of coincidental overlaps in shortest path profiles between non-isomorphic environments do not undermine the overall trend, as the majority of pairs exhibit clear distinctions.

### Zero-shot navigation across environments with multiple subgoals

Next, we examined how TAG’s topology-aware exploration could enhance goal-directed learning, particularly in complex scenarios involving multiple subgoals. While random walk maps have proven effective for single-goal navigation tasks^11^, their efficacy in multi-subgoal navigation remains largely unexplored. Such scenarios present a unique challenge, as they require the seamless integration of multiple policies within a unified task framework. Although Successor Features Q-Learning (SFQL)^9^ addresses this challenge using multiple successor features (SFs), it is not biologically plausible and relies on multiple separate representations for task completion. To resolve these issues, we implemented a novel zero-shot navigation strategy that leverages topological information of our TAG.

Our TAG-based navigation strategy exploits the fact that topology inherently defines the sequential order of nodes required to reach a specific goal (Fig. 7a). Using this topological structure, the agent navigates subgoals in the order required to reach the goal in the topology graph. For cases where multiple subgoals are associated with the same topological node or edge, the agent employs a greedy strategy, visiting subgoals in order of proximity. If multiple distinct paths lead to the goal and multiple subgoals lie along the same topological sequence, the agent randomly selects one topology node to visit first and proceeds to the next.

**Figure 7.**
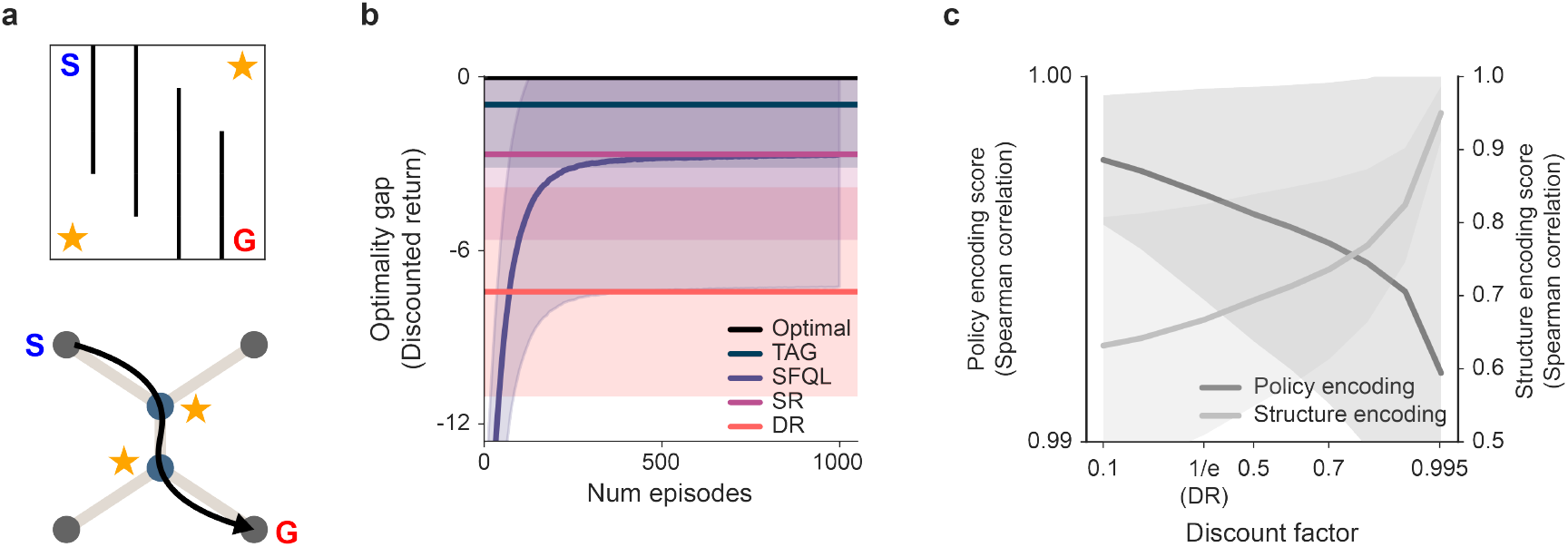
Topological subgoal navigation. **a**, An overview of the decision-making logic for navigation using topology. The agent determines the sequential order of subgoals based on the topological structure of the environment. Subgoals are visited in the order defined by the topology graph. For subgoals associated with the same topological node or edge, the agent adopts a greedy approach, visiting them in order of proximity. In cases where multiple paths lead to the goal and subgoals lie along the same topological sequence, the agent randomly selects one node to visit first before proceeding further. **b**, The agents were tested on the randomly generated environmental structure and reward locations, where four subgoals and one goal yielded rewards of 3 and 10, respectively. The optimality gap is calculated as the difference between the discounted returns of the model’s performance and the optimal trajectory obtained using dynamic programming. Discounted returns are computed with *γ* = 0.9999, to prevent too much penalty being applied when deviating from the optimal trajectory. **c**, Comparison of policy encoding score and structure encoding score for various discount factors. Scores are computed by calculating the Spearman correlation between the model’s representations and the optimal geodesic distances. Shaded areas represent standard deviations across 1000 randomly generated mazes.

The TAG model was pitted against three baselines: SFQL, DR, and SR, focusing on multi-subgoal navigation performance. SFQL was trained over 1000 episodes, and its final performance was compared with TAG, DR, and SR, which assumed knowledge of the exact environment structure. For DR, the subgoal rewards were incorporated into the nonterminal states (refer to Supplementary Section 4 and 7 for more discussions on utilizing DR for subgoal navigation), and for SR, rewards for both subgoals and goals were embedded in the reward vector (Methods). Dynamic programming was used to calculate the optimal trajectory and discounted returns as a benchmark.

The TAG model achieved near-optimal performance using zero-shot planning, significantly outperforming other models(Fig. 7b). While SFQL successfully navigated toward a goal and visited all subgoals, it often used suboptimal trajectories between subgoals, resulting in greater discounted reward losses compared to the optimal trajectory. One striking result is that SR outperformed DR. Despite the diffusion of rewards across the SR placefield, which created multiple local maxima, some of these maxima directed the agent toward subgoals. However, SR often missed certain subgoals while heading toward the goal. In contrast, DR’s placefield was less effective in guiding the agent to visit subgoals. Paradoxically, this limitation stems from DR’s optimization for reaching the goal state.

DR’s default configuration, with *r*_*N*_ = −1 and *µ* = 1, is functionally equivalent to a random-walk SR with a discount factor of *γ* = 1*/e*. Specifically,

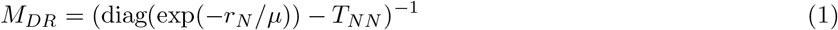

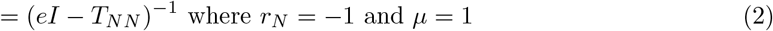

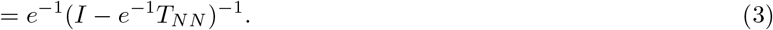

This low discount factor penalizes the transition matrix power with high *t* more than high discount factors, thereby focusing the agent on the shortest trajectories to the goal. This ensures optimal policy encoding. However, in most cases, the subgoals are located outside of the optimal trajectory, resulting in *t* being high. Although the trajectories with high *r*_*N*_ should become higher, the exponential decay of the discount factor easily suppresses the increases in *r*_*N*_. Moreover, the fact that *r*_*N*_ is bounded by zero makes it even difficult to deviate from the optimal trajectory.

We found that, while the low discount factor is beneficial when there is one goal^11^, the high discount factor enhances the structure encoding in the successor coordinate (Fig. 7c). This is because higher discount factors assign greater weight to eigenvectors crucial for encoding structural information. Notably, TAG demonstrates the ability to effectively reconcile these competing requirements by maintaining a proper balance between the low and high discount factors.

### Scalable, biologically plausible neuromorphic computing

We demonstrated that TAG can be implemented within a transformer-like architecture, building upon recent work showing how transformers can compute place cells from grid cells^48,49^. This translation is critical, as it bridges biological theory with scalable machine learning models. This architecture not only accounts for the necessity of multiple heads and stacks to support complex computations—enabling distinct functional roles across the architecture—while extending these computational principles to decision-making. Notably, this defines a new, scalable building block for topology-aware deep learning.

Our architecture employs eigenvectors of SR as positional encodings, a method analogous to transformer-based graph neural networks^50,51^which utilize eigenvectors of the graph Laplacian for similar purposes. By employing a shared positional encoding and varying the weights, the architecture ensures that different heads serve distinctly different computational roles from the same input data (refer to Supplementary Fig. 1).

In the policy head, the model calculates the SR matrix M using self-attention between grid codes G, weighted by the square root of SR eigenvalues with a low discount factor,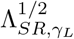. As the value input, features Φ encode next goal that the agent targets at the moment, while *w* represents their corresponding rewards. The output of this head is a place code *P*, which guides decision-making by determining the agent’s immediate navigation targets.

The structure head reconstructs the transition matrix *T* from the grid codes 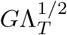 through eigende-composition. A linear readout *W*_readout,*B*_ is then applied to derive a boundary code B from the transition matrix. Subsequently, self-attention on *T* produces a squared transition matrix *T* ^2^, capturing second-order transition relationships. This is further processed using a value readout *W*_readout,*C*_ to compute the corner code *C*.

In the distance head, grid codes weighted by the square root of SR eigenvalues with a high discount factor, 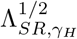, are processed using distance-based self-attention^52,53^. This mechanism measures state dissimilarities in the successor coordinate, enabling the generation of a topology graph. The edge weights of this graph are determined by the output of the distance head, which incorporates distances between the nodes into the graph.

Overall, the structure head contributes to the generation of the topology graph, while the distance head determines its edge weights. This topological structure of the environment defines Φ in the policy head, a variable that determines which subgoal to navigate next. By combining these components, the architecture achieves efficient topology-based decision-making, leveraging shared positional encodings to maintain consistency across the heads while enabling specialized computations.

## Discussion

This study underscores the critical role of topology in enabling structure and reward generalization, emphasizing the role of policy-independent grid codes in representing environmental topological structures while reflecting the geodesic distance between the topological state labels. We proposed a computational framework called TAG, integrating grid, place, boundary, and corner codes into a unified hierarchy double-loop architecture reminiscent of the 0–1–2-dimensional features counted by Euler characteristic^36^, elucidating the distinct contributions of each neural code in understanding topology and facilitating decision-making. The inner loop of TAG composed of place and boundary codes assisted in decision-making and rapid, policy-independent environmental learning by calculating structural SPE. The outer loop of TAG consisting of boundary and corner codes contributed to encoding topological features. This double loop architecture yielded a topology-aware grid representation. We demonstrated TAG’s biological plausibility across three key findings: rapid, boundary code-driven grid-scale adaptation following environmental resizing; accurate replication of CA1 and subicular topology-encoding manifolds and their relative dimensionalities; and the essential contribution of low-eigenvalue (WS) versus high-eigenvalue (SS) grid components to robust topological embedding.

The TAG grid code effectively showed robustness against topology-preserving environment augmentations, defined as noise, shift, scale, and rotation. This robustness enabled TAG to zero-shot transfer across isomorphic environments, outperforming the compositional generalization method. At the same time, TAG maintained its capacity to distinguish between non-isomorphic environments. This allowed TAG to learn novel isomorphic environmental structures more quickly than topology-agnostic exploration strategies. TAG employed dual discount factors, prioritizing structure and policy encoding performances respectively, to perform nearly optimal zero-shot navigation in complex scenarios requiring planning through multiple subgoals. Lastly, TAG shows the potential for extending the framework to deep learning models through a transformer-based analogy, proposing a building block for topology-aware deep learning models.

Beyond cognitive maps, neuroscience has uncovered cognitive graphs—abstract, graph-like representations of state relationships that need not be purely spatial^54–58^. TAG offers a mechanistic account of how such graphs might be constructed neurally: by combining policy-independent grid, place, boundary and corner codes into an explicit topology graph, it shows how the brain could transform raw spatial inputs into an abstract graph representation for flexible planning and generalization.

While TAG’s boundary and corner codes are used to capture the environmental topology, these neurons also encode environment-specific geometric details. Environmental topology can remain the same while the physical shapes of environments differ, leading to variations in boundary and corner codes. Some prominent examples are seen in^59,60^, where environments with different shapes, sharing identical topology, are distinguishable based on neural codes. This suggests that while certain boundary and corner codes contribute to topology encoding, others may encode shape-specific information. Distinguishing which boundary and corner codes are used for topology encoding versus those representing environment-specific details could offer intriguing avenues for further exploration.

As discussed, our model aligns with transformer-based architectures by incorporating multi-head and stacked structures. While our framework employs three heads and at most two stacks, transformer-based graph neural networks typically utilize more extensive head and stack configurations. Exploring how additional heads or stacks could enhance the model’s capacity to process more complex environmental features would be an interesting direction. Additionally, the transformer analogy might guide the development of deep learning models that extend our framework, a potential future endeavor.

While our findings advance our understanding of hippocampal information processing, several limitations remain that future work could address. First, the current definitions of boundary and corner codes rely on discrete state representations, making them incompatible with continuous state spaces. Extending our model, which heavily depends on matrix operations, to continuous state spaces poses challenges. Referencing methods like deep reinforcement learning for proto-value function learning in continuous state spaces^61^ could offer viable pathways for such expansions.

Second, although TAG assumes a perfectly policy-independent grid code, real grid cells do show gradual reward biases when a goal remains fixed for long periods^62^. Those slow-timescale distortions likely arise from a form of cumulative policy regularization—quite distinct from the robust grid coding of TAG under changing reward. Incorporating a complementary long-horizon grid code update mechanism that gradually integrates enduring reward signals would be an interesting direction for future work.

Third, topology is not a universal descriptor of environmental structure. For instance, environments with distinct room-like configurations, such as the four-room setup, are encoded as torus-like topologies rather than room-specific nodes. Although clustering SR row vectors based on stationary distributions or using normalized cuts^46^ might resolve these ambiguities, such methods lie beyond the scope of topology encoding. Moreover, the thinning algorithm, used for extracting topology, lacks robustness in some cases and introduces instability. Addressing these algorithmic limitations would improve the reliability of topological inference.

Our study provides a theoretical foundation for topology-based navigation and environmental encoding. While addressing the discussed limitations requires further research, our findings pave the way for applying topology-driven models to continuous spaces and reward-sensitive tasks. Future work should explore these directions, potentially bridging gaps between biological and computational models of spatial cognition.

## Methods

### Maze environments

The environment consists of a 10 × 10 grid world with four available actions: left, right, up, and down. Each environment is randomly generated while controlling the proportion of blocked states within the grid. The proportion of blocked cells is sampled from ten evenly spaced bins ranging from 0 to 0.5. For each bin, 100 unique mazes are randomly generated, resulting in a total of 1,000 environments. To ensure navigability, each generated maze undergoes a connectivity validation process using the Breadth-First Search (BFS) algorithm, and only environments where all non-block states are fully connected are retained. To prevent environments with a corresponding topology graph consisting of only a single node from trivially influencing the simulation results, we excluded such cases in simulations utilizing TAG’s topology-awareness (Fig. 4–7). In these cases, all environments are inherently isomorphic, and both grid code and environment distance measurements would trivially yield zero, potentially overstating robustness to isomorphic transformations while simultaneously exaggerating sensitivity to non-isomorphic variations. Otherwise, all 1,000 generated environments were used in the simulations.

### Topological state abstraction

We classify states into three labels: junctions, edges, and dead-ends. Junctions are states connected to other junctions or dead-ends, requiring the agent to make a navigational choice. Dead-ends are terminal states that the agent should generally avoid unless they are rewarding or specified as goals. Edges act as connectors, linking junctions to other junctions, dead-ends to other dead-ends, or junctions to dead-ends. This topological abstraction simplifies the state space while retaining essential features critical for effective navigation and generalization. However, the resulting topological state abstraction may vary depending on the perspective chosen, and multiple valid abstractions could exist for a single environment. This ambiguity is particularly pronounced in environments with fewer obstacles, where insufficient regularization constrains the assignment of topological state labels.

To ensure consistency and reliability in determining state labels, we utilize the Zhang-Suen thinning algorithm^63^ on the maze image. This iterative algorithm removes non-essential non-block states, leaving a skeletal representation that captures the maze’s fundamental topology. Such skeletal representations are widely used to extract structural features from complex objects^64^.

The preprocessing involves drawing black walls between unconnected states and marking states entirely surrounded by walls as black. The maze’s size is tripled to maintain a minimal pixel count while ensuring each state occupies an odd number of rows and columns for thinning unbiased towards either direction. This adjustment prevents skeleton lines from being biased to one side and creates sufficient space for walls.

During iterative thinning, only skeletal pixels remain, representing the maze’s structural connections. Connections for each pixel are examined in eight directions: first the cardinal directions (up, down, left, right), followed by the diagonals (top-right, top-left, bottom-right, bottom-left). These skeletal pixel nodes are classified based on the number of connections: nodes with three or more connections are labeled as junctions, nodes with one connection as dead-ends, and nodes with exactly two connections as edges. Each state is assigned the label of its nearest skeletal pixel node, determined using Euclidean distance.

For the final graph representation (e.g. Fig. 1c), pixels with exactly two connected edges are removed to simplify the topology graph and ensure accurate edge representation. This process provides a robust framework for classifying states and constructing a meaningful abstraction of the environment’s topological structure.

### Successor distance and diffusion distance

Inspired by the diffusion coordinate^65^, we define the successor distance:

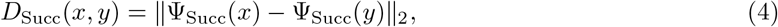

which is the Euclidean distance in the successor coordinate^66^. The successor coordinate is defined as:

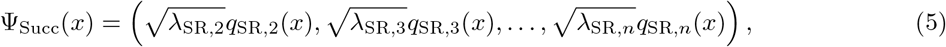

where *λ*_SR,*i*_ and *q*_SR,*i*_ represent the *i*-th eigenvalue and eigenvector of the SR matrix, respectively.

Notably, the first eigenvector and eigenvalue are omitted because the first eigenvector assigns uniform values across non-block states^67^. As a result, it does not contribute to distinguishing between states when calculating the Euclidean distance in the successor coordinate.

The successor distance is analogous to the diffusion distance^65^, defined as:

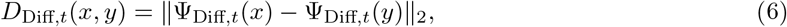

where the diffusion coordinate is defined as:

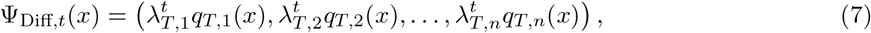

with *λ*_*T,i*_ and *q*_*T,i*_ defined as the *i*-th eigenvalue and eigenvector of the transition matrix, and *t* as a scale parameter.

Both metrics reflect Markov chain structures, but the successor distance directly utilizes the SR eigenpairs and omits the scale parameter. Note that full grid cell activation was used, rather than restricting to the non-negative components (see Supplementary Note 8 for details).

### Gromov-Wasserstein distance between grid codes

The distances between the grid codes were assessed using the Gromov-Wasserstein (GW) distance^68^, which measures the loss between the model’s grid code and the ground-truth grid code derived from random-walk SR. The GW distance quantifies discrepancies between pairwise distances in two metric spaces, effectively capturing the structural alignment between the grid codes. By focusing specifically on the successor distances between junction and dead-end topology labels, rather than all state pairs, we evaluate how accurately the model captures the critical topological relationships. This focus is justified because junctions and dead-ends are key structural elements that define the global topology of the environment, making them particularly relevant for assessing topology embedding accuracy. When comparing grid codes between different environments (e.g. original and augmented environments at Fig. 4, or between non-isomorphic environments at Fig. 6), the GW distance may capture the differences between the absolute successor distance between the corresponding topological edges. Therefore, we calculated GW distance after normalizing the size of both grid codes. Importantly, the GW distance is invariant to manifold rotation, requiring only scale normalization to measure topological discrepancies accurately. The discount factor was set as *γ* = 0.995, unless otherwise specified. Except for the simulations in Fig. 1, we used 10-dimensional successor coordinates to ensure that the total dimensions match between the environments with different sizes, while considering all the dimensions with nontrivial eigenvalues of SR (see Fig. 2 for the distribution of SR eigenvalues with *γ* = 0.995).

### Gromov-Hausdorff distance between the environment structures

To quantify environmental structural similarity, we adopt the Gromov-Hausdorff (GH) distance^69^, a metric designed to compare topologies. The GH distance quantifies the maximal discrepancy in shortest path distances between nodes after an optimal alignment, effectively capturing the overall global structure of a topology graph. By focusing on the overall geometry rather than requiring an exact match of local edge weights, GH distance preserves the ranking of shortest path distances, enabling robust comparisons between environment topologies and reliably distinguishing between topologically isomorphic and non-isomorphic environments.

### Topology label classification using transformer classifier

The input to the model consists of a 32-dimensional feature vector that encodes various structural and positional information about the maze. Corner information is represented using 8 binary dimensions, indicating the four cardinal directions (north, east, south, west) and whether each corner is concave or convex. Boundary information is captured in 4 binary dimensions, corresponding to the presence or absence of boundaries in the four cardinal directions. The place code, which encodes the agent’s positional information, is initially represented as a 100-dimensional vector for a 10 × 10 maze and is reduced to 10 dimensions using a fully connected network, with all values being positive. Grid code information is represented by 9 dimensions, corresponding to the 9 largest eigenvalues (excluding the first) of the successor representation, starting from the Fiedler vector. The grid code values are continuous and range between -1 and 1. Additionally, each state includes a block code represented by a single binary dimension, indicating whether the state is blocked (0 or 1). These features are concatenated with a 32-dimensional 2D sinusoidal positional encoding^70^ to form a 64-dimensional input to the transformer. For ablated inputs, the respective dimensions (e.g., corner, boundary, place, or grid code) were filled with zeros to simulate their absence during training and evaluation.

The transformer model comprises six stacked layers with eight attention heads, as described in the original Vaswani architecture. It takes the 64-dimensional combined input (32-dimensional input feature vector and 32-dimensional positional encoding) and processes it to predict maze topology labels. The model is trained using cross-entropy loss between the predicted and ground-truth labels, with the final labels determined via argmax over the output probabilities. Training was conducted on 1,000 randomly generated mazes, and the model was evaluated on 200 unseen mazes. Ground truth labels for maze topology were generated using the Zhang-Suen thinning algorithm, which extracts the skeletal structure of the environment.

### Experimental details

#### Exploration policy entropy analysis

In the experiment testing the grid code training performance (Fig. 2b), we modified the exploration policy entropy as follows: H0 is where the up, down, left, and right have equal probability of 0.25. On the other hand, H4 is where up and left have probability of 0.05, and right and down have 0.45. The probability for up and left are diminished by 0.05, and right and down are increased by 0.05 as the entropy level is increased by 1.

#### Measuring the contributions of neural codes

We used Shapley value^71^ to quantify the contribution of each neural code.

#### Aligning grid codes between environmental topology-preserving augmentations

We used the Procrustes algorithm^72,73^ to align the set of TAG grid codes of the augmented environment with that of the original environment. This was applied along with the manifold size normalization.

#### Measuring the graph similarity

We used graph edit distance^74^ to measure the structural similarity between two graphs, focusing only on their connectivities, not edge weights.

#### Barry et al., 2007

We instantiated a 10 × 10 discrete maze and generated four environment geometries by blocking sets of cells following Barry et al. (2007): 1. Large square: no blocked cells (10 × 10 open). 2. Vertical rectangle: right quadrants blocked, yielding a 10 × 7 open region in the left half. 3. Horizontal rectangle: lower quadrants blocked, yielding a 7 × 10 open region in the upper half. 4. Small square: lower-right, lower-left, and upper-right quadrants blocked, yielding a contiguous 7 × 7 open region in the top-left. For each geometry, an unbiased random-walk policy was executed for 20,000 time steps. After learning, two-dimensional spatial autocorrelograms of the TAG grid codes were computed. To quantify grid rescaling, we searched over horizontal and vertical scale factors to maximize the Pearson correlation between the baseline and each probe-geometry autocorrelogram.

#### Nakai et al., 2024

We implemented a Double D maze implemented as a 45 × 45 discrete grid. To compare model and neural manifold dimensionality, we estimated intrinsic dimensionality using the Grassberger–Procaccia algorithm^75^ on successor-coordinate embeddings derived from 1,000 randomly generated 10 × 10 mazes. To reflect the real neural activity that has response field larger than one state, we applied an exponential decay 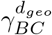 where *γ*_*BC*_ is a decay parameter and *d*_*geo*_ is a geodesic distance. To better match BVC tuning^20,23^, the TAG boundary code was extended beyond immediate (1-step) wall distances by computing activations at multiple distances from the nearest wall. This adjustment reflects BVC receptive fields that respond to specific distances to environmental boundaries. Because biological corner cells exhibit responses to multiple corners^21^, TAG corner codes were generated by combining multiple corners of the same type (either convex or concave) via a random weighted sum.

#### Guo et al., 2024

To emulate the differing learning stages observed in Guo et al., the TAG model’s transition matrix was randomly sparsified with probabilities of 0.20 for the early training stage and 0.10 for the late training stage. This procedure was repeated for eight different random seeds. A zero entry denotes an unlearned (though physically available) state transition; the remaining transitions in each row were renormalized to maintain a uniform random-walk policy despite partial knowledge. To classify SS versus WS place codes, SR eigenvalues were sorted and a cutoff set at the 230th eigenvalue: eigenvectors 1–230 were labeled SS, and eigenvectors 231 onward labeled WS. The environment was implemented with a 45 × 45 grid world environment.

### Topology-preserving environment augmentations

We define four topology-preserving environment augmentations: noise, shift, scale, and rotation (Fig. 4a). Noise augmentation introduces unstructured perturbations by randomly adding blocks at various positions within the environment. As the most challenging scenario for generalization, noise augmentation disrupts the local state-transition dynamics while preserving the overall topology of the environment. This tests the grid code’s ability to maintain structural relationships under irregular and unpredictable modifications. In shift augmentation, the block states are translated by *x* units along the row direction and *y* units along the column direction. In cases where the shifted environment causes the length of paths to increase, edge lengths may grow while the topology remains unchanged. To prevent additional paths from altering the topology, blocks are added to newly generated states, extending the boundaries of the environment and preserving its structural integrity. Scale augmentation increases the size of the environment by dividing each state into *k*^2^ smaller states based on a given scale factor *k*. While the proportions between states are maintained, the absolute distances and resolution of the grid code representation change. This requires normalization to ensure that topological relationships are consistently preserved despite these alterations. Rotation augmentation rotates all block states counterclockwise *t* times, determined by a rotation factor *t*. As a control condition, rotation affects only the indexing of states without altering the underlying state-transition dynamics or topology. Consequently, the TAG grid code is expected to remain entirely invariant under this augmentation, demonstrating robustness to transformations that preserve all structural and relational aspects of the environment.

### Topology-aware exploration strategy

For topology-aware exploration, the learning agent must focus on exploring new edges rather than exhaustively mapping the precise structural details. The agent considers two hypothetical scenarios for each unexplored state-transition: one where the transition is assumed possible and another where it is assumed impossible. Using these assumptions, the agent generates two distinct topology graphs that represent the potential structural outcomes. By comparing these graphs, the agent identifies junctions or edge states that are most likely to yield new topological edges. For each identified candidate, the agent constructs all possible topology graphs by considering different combinations of potential new edges. To identify the best-matching candidate topology graph to the topological prior, the agent computes the GW distance between each candidate graph and the prior topology graph. The agent then selects the nearest unexplored state-transition corresponding to the most likely new edge predicted by the best-matching candidate topology graph and prioritizes it for exploration. By iteratively refining its understanding of the environment through this systematic process, the agent ensures efficient and topologically informed exploration. Refer to the Supplementary Note 3 for full details.

### Baselines for subgoal simulation

To evaluate the effectiveness of the TAG algorithm, we compare its performance against three baselines: SFQL^9^, DR^11^, and SR^25^. The SFQL baseline is trained over 1000 episodes, and its final performance is assessed. For the DR baseline, the agent is assumed to have precise knowledge of the environment’s structure. In this representation, terminal and nonterminal states are treated separately. Goal states are classified as terminal, with rewards exponentially weighted and applied at the end of navigation. Subgoal states are treated as nonterminal and incorporated into the computation of the DR matrix. Considering the invertibility and characteristics of DR matrix, *r*_*N*_ must be bounded by zero:

#### Theorem 1

*Let n* ∈ ℕ *and µ >* 0. *Suppose R* = diag(*d*_1_, *d*_2_, …, *d*_*n*_), *where d*_*i*_ = exp(*r*_*i*_*/µ*), *and let T be an n* × *n row-stochastic transition matrix. If r*_*i*_ *<* 0 *for all i, then the matrix M* = *R*^−1^ − *T is invertible*.

(refer to Supplementary Note 4 for proof and more details).

Since subgoals may have positive rewards or cases where no subgoals exist, we apply an activation function:

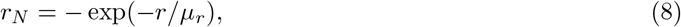

where *r* is the environmental reward, and *µ*_*r*_ is a temperature parameter. This ensures *r*_*N*_ increases as *r* increases while remaining bounded by 0. After the agent visits a subgoal, the corresponding element of *r*_*N*_ is set to 0, reflecting its completion.

The SR baseline follows a similar structure, with the state values computed using:

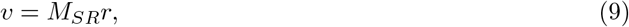

where *M*_*SR*_ is the SR matrix, and *r* includes both goal and subgoal rewards. As with DR, the reward for a visited subgoal is set to 0 once completed. For all models, we calculate the discounted returns to assess navigation performance. The discounted sum of returns allows us to measure not only whether the agent collects all subgoals but also whether it visits them in an optimal order. Optimal performance is derived using dynamic programming, which provides the benchmark trajectory and discounted returns for comparison.

## Acknowledgements

We thank Dongqi Han, Yoondo Sung, and Minsu Abel Yang for their insightful comments and feedback. This work was supported by Electronics and Telecommunications Research Institute(ETRI) grant funded by the Korean government (25ZB1100, Core Technology Research for Self-Improving Integrated Artificial Intelligence System), Institute of Information & communications Technology Planning & Evaluation (IITP) grant funded by the Korea government (MSIT) (No. RS-2023-00233251, System3 reinforcement learning with high-level brain functions; No. RS-2019-II190075 Artificial Intelligence Graduate School Program(KAIST)), the National Research Foundation of Korea(NRF) funded by the Korean government (MSIT) (No. RS-2024-00439903, RS-2024-00341805), the MSIT(Ministry of Science, ICT), Korea, under the Global Research Support Program in the Digital Field program(RS-2024-00436680) supervised by the IITP(Institute for Information & Communications Technology Planning & Evaluation).

## Author contributions

H.K. and S.W.L. conceived and designed the study. H.K. implemented the core TAG model and ran the simulations. D.C.M. and N.J. contributed to model development and conducted comparative analyses. H.K. and S.W.L. wrote the paper. H.K., D.C.M., and S.W.L. reviewed and edited the paper. All authors approved the final version for submission.

## Supplementary

### 1 Relationship between eigenvectors of SR, transition matrix, and graph Laplacian

The grid code of the SR model (*M*), which is the eigenvectors of the SR matrix (*Q*_*SR*_), are identical to those of the transition matrix (*Q*_*T*_) and closely related to the eigenvectors of the normalized graph Laplacian (*Q*_L_)^45,76^:

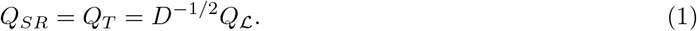

Specifically,

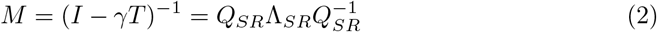

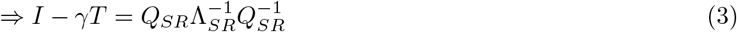

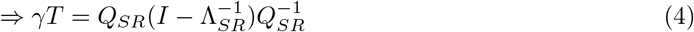

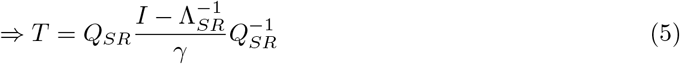

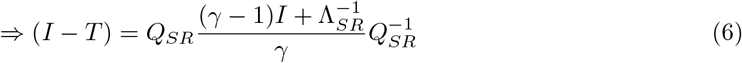

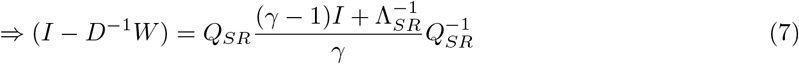

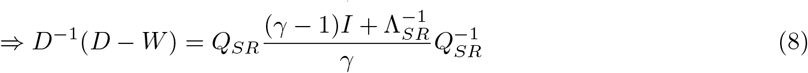

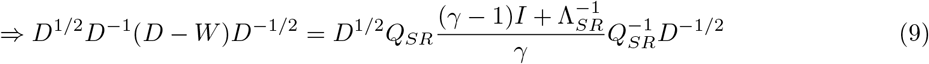

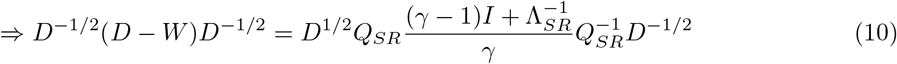

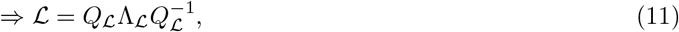

where *D* is the degree matrix and *W* is the weight matrix of the graph Laplacian.

Since *T* = *D*^−1^*W*, it follows that all elements in *W* must be uniform in a policy-independent Markov chain scenario, where each state transitions uniformly to its neighbors.

### 2 Pseudocode for policy-independent grid code learning

#### Algorithm 1

UpdateSR

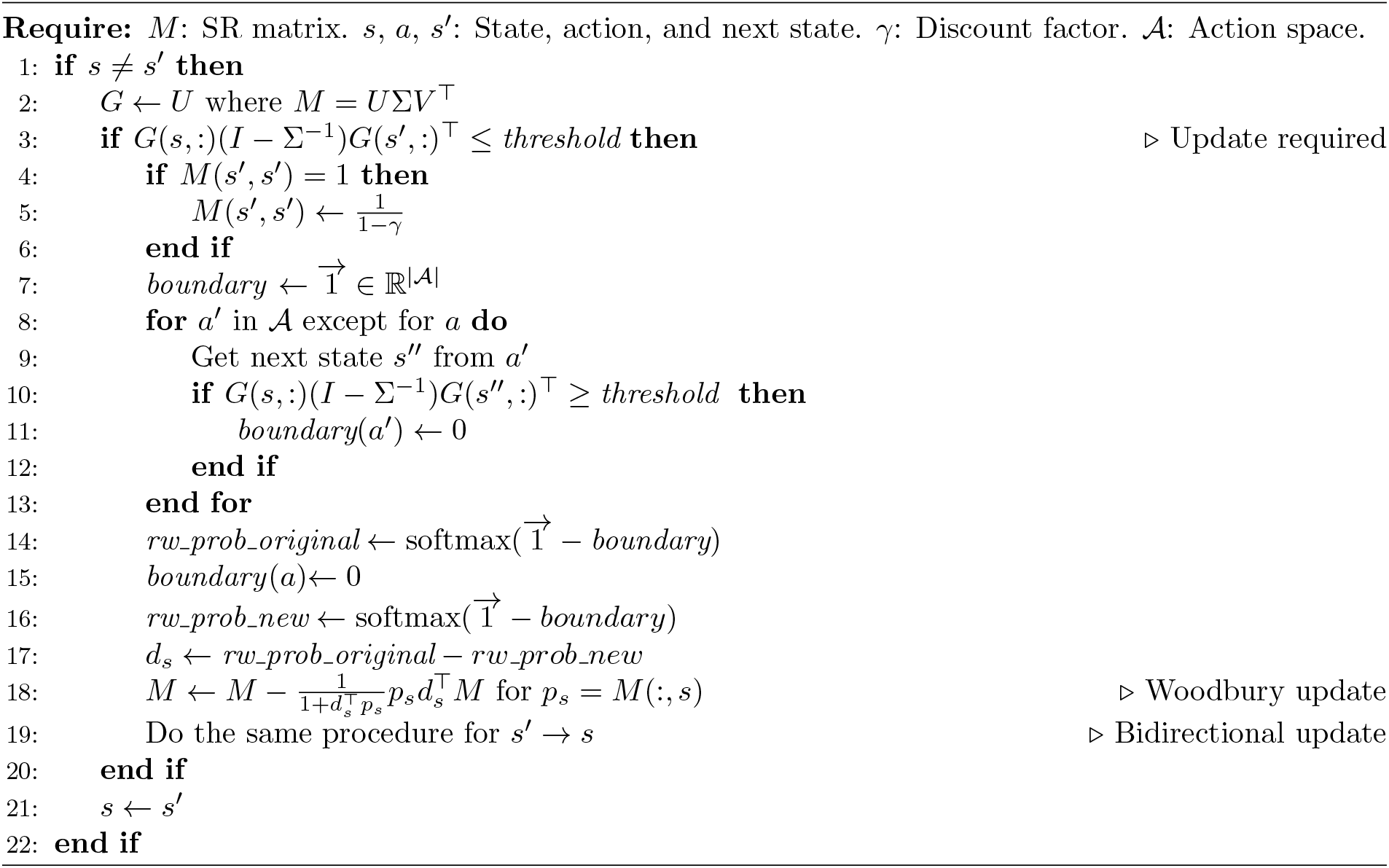

Note that the updates occur bidirectionally (*s*_*t*_ → *s*_*t*+1_ and *s*_*t*+1_ → *s*_*t*_), following Keck et al.^77^. This approach leads to sample-efficient learning and faster adaptation to environmental structure with fewer exploratory steps. The agent initially assumes all state transitions are impossible, progressively identifying feasible transitions through exploration.

Here, we use singular value decomposition (SVD) instead of eigendecomposition due to the numerical instability associated with the latter. While eigendecomposition is generally sufficient for analyzing the characteristics of eigenvectors, its use in iterative algorithms can lead to the accumulation of numerical errors. In contrast, SVD provides greater numerical stability in such contexts. Additionally, it has been shown that singular vectors effectively approximate proto-value functions, generalizing eigendecomposition^45,76,78–80^.

Below is the justification of setting *M* (*s, s*) as 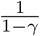:

**Theorem 2**. ^*81*^ *The sum of the SR’s row vector for every approachable state is* 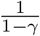.

*Proof*. For every approachable state *s*, we have:

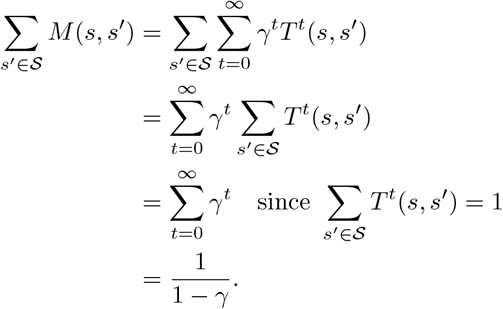

### 3 Pseudocode for the TAG exploration algorithm

#### Algorithm 2

TAGExplore

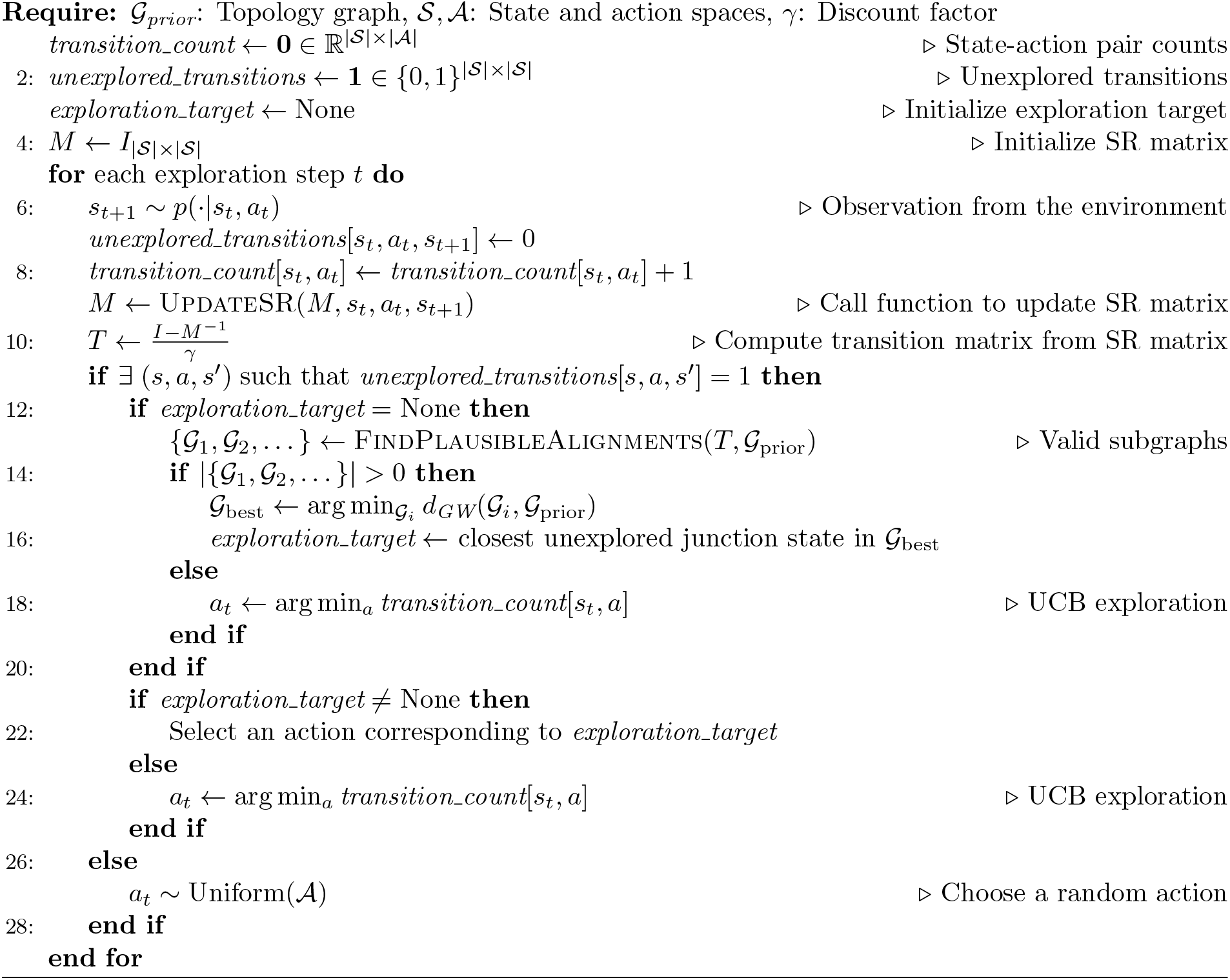

#### Algorithm 3

Find Plausible Alignments

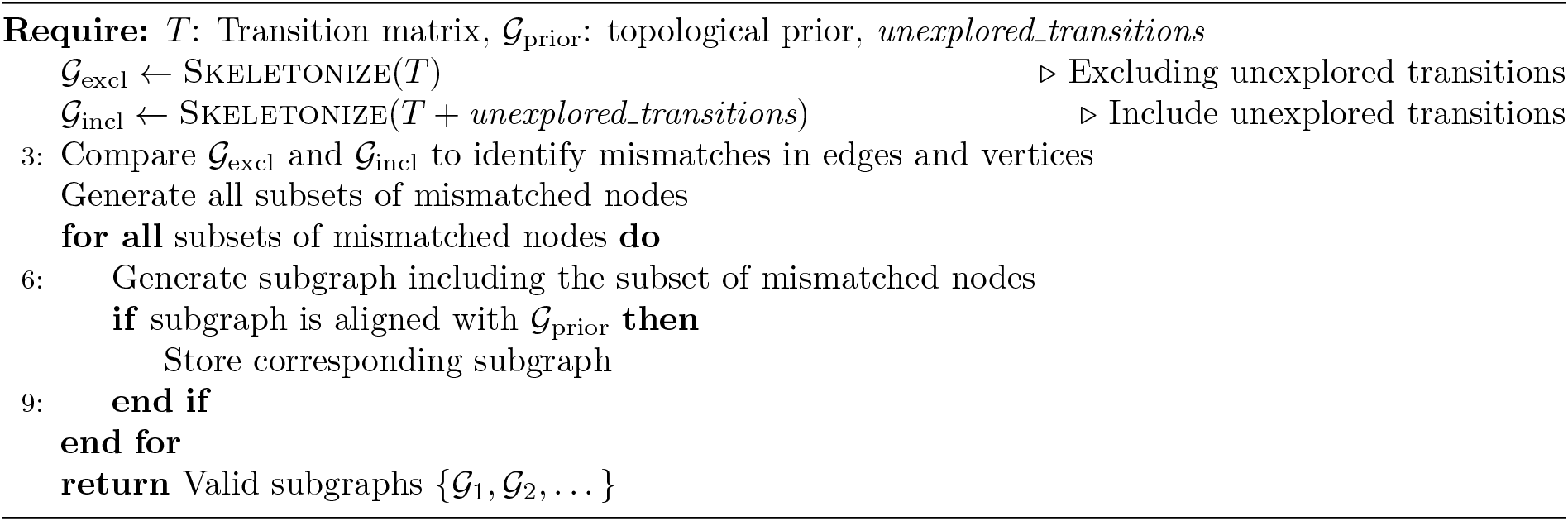

Skeletonization refers to the Zhang-Suen thinning algorithm described in the Methods.

### 4 A constraint for the nonterminal rewards of DR

In this section, we justify our choice of activation function applied to the nonterminal rewards of default representation (DR) for subgoal navigation: *r*_*N*_ = − exp(−*r/µ*_*r*_). First, we argue that *r*_*N*_ *<* 0 ∀*r*_*N*_ is a sufficient condition for the invertibility of the DR matrix. Subsequently, we demonstrate that non-negative *r*_*N*_ may cause issues in terms of the invertibility of the DR matrix, its interpretation in a reinforcement learning context, and its effectiveness in supporting spatial navigation.

#### Definition

The DR matrix *M* is expressed as:

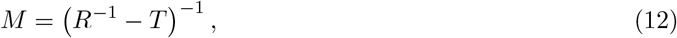

where 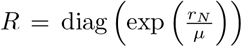, while *r*_*N*_ represents rewards at nonterminal states, *µ* represents a temperature parameter. Note that we used notation *µ* instead of *λ* originally used by the authors to avoid confusion with eigenvalues.

The value function of DR is given by:

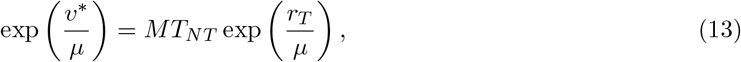

where *v*^∗^ is the value vector, *r*_*T*_ represents rewards at terminal states, *T*_*NN*_ is the transition matrix between nonterminal states, and *T*_*NT*_ is the transition matrix from nonterminal to terminal states.

#### Invertibility of DR

*Proof of Theorem 1*. Since *R* is invertible, we have *R*^−1^ − *T* = *R*^−1^(*I* − *RT*). Noting that det(*R*^−1^ − *T*) = det(*R*^−1^) det(*I* − *RT*) and that det(*R*^−1^)/= 0, it follows that *R*^−1^ − *T* is invertible if and only if *I* − *RT* is invertible.

Recall that a square matrix *A* is invertible if and only if 0 ∈*/σ*(*A*) (where *σ*(*A*) denotes the spectrum of *A*). In particular, for *A* = *I* − *RT*, we have 0 ∈ *σ*(*I* − *RT*) precisely when there exists a nonzero vector *v* such that (*I* − *RT*)*v* = 0; that is, when *RTv* = *v*, which is equivalent to 1 ∈ *σ*(*RT*). Thus, *I* − *RT* is invertible if and only if 1 ∈*/σ*(*RT*).

Since *T* is row-stochastic, the sum of the entries in the *i*th row of *RT* is 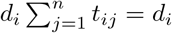. Hence, the ∞-norm of *RT* is

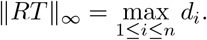

Because the spectral radius *ρ*(*B*) of any matrix *B* satisfies *ρ*(*B*) ≤ ∥*B*∥_∞_, we obtain the key inequality

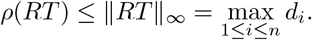

If *r*_*i*_ *<* 0 for all *i*, then *d*_*i*_ = exp(*r*_*i*_*/µ*) *<* 1 for each *i*; hence, max_1≤*i*≤*n*_ *d*_*i*_ *<* 1 and *ρ*(*RT*) *<* 1. This implies every eigenvalue *λ* of *RT* satisfies |*λ*| *<* 1, in particular, 1 ∈*/σ*(*RT*). Consequently, *I* − *RT* (and hence *R*^−1^ − *T*) is invertible.

Although having some *r*_*N*_ equal to zero or positive does not necessarily prevent the DR matrix from being invertible, there are notable issues when using *r*_*N*_ = *r*.

1. **When all the nonterminal rewards are zero:** This is inherently equivalent to

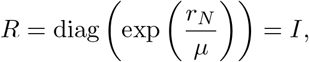

where *I* is the identity matrix. As stated in the proof above, *I* − *T* is invertible if and only if 1 ∈*/σ*(*T*). However, every transition matrix has an eigenvalue equal to 1^65^. Therefore, *I* − *T* is not invertible, meaning that the DR matrix cannot be computed in this case.
2. **When** *R* = *dI* **with** 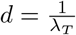 **for some eigenvalue** *λ*_*T*_ **of the transition matrix:** This represents an extreme edge case, but in this situation, there exists an eigenvalue of *RT* equal to 1, making *I* − *RT* non-invertible.

Moreover, there can be additional cases that result in 1 ∈ *σ*(*RT*). To avoid all such scenarios, it is safer to enforce the condition *r*_*N*_ *<* 0.

#### State-wise discounting of the rewards

We discuss the meaning of *R* in a reinforcement learning context, arguing that the diagonals of *R*, denoted as *d*_*i*_, represent state-wise discount factors. Hence, *r*_*N*_ should be bounded by zero to ensure that the discount factors lie within the range (0, 1).

To show this, we rewrite the DR matrix formula:

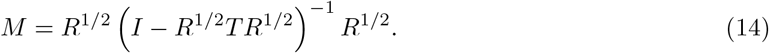

This formulation shows that each transition matrix element *t*_*ij*_ is discounted by the geometric mean of the respective state discount factors:

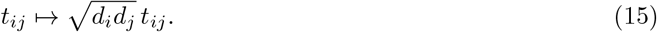

As transitions propagate, each intermediate state in the path introduces additional discounting. For example, for (*R*^1*/*2^*TR*^1*/*2^)^2^, the element becomes:

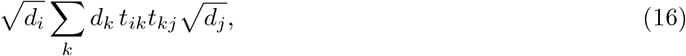

where intermediate states also contribute their respective discount factors. The final discount applied to a path reflects the product of the discount factors of all states involved, including the starting, intermediate, and terminal states.

This discounting behavior leads to the diffusion of state-wise penalties, as the discount factors affect not only the state itself but also its neighbors through transition dynamics.

However, with nonnegative values of *r*_*N*_, the corresponding *d*_*i*_ would be 1 or greater than 1. This does not align with the reinforcement learning framework, where the discount factor should be strictly less than 1 while remaining positive.

#### Navigation performance

Positive *r*_*N*_ values also affect navigation performance. SR-related models, including DR, guide navigation toward directions that yield higher place field values. However, if *r*_*N*_ becomes positive for some states, the largest eigenvalue of *RT* may exceed 1. This results in the corresponding eigenvalue of the DR matrix becoming negative with a large magnitude, as the eigenvalue of DR is given by:

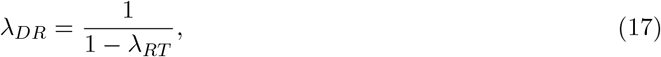

where *λ*_*RT*_ is the eigenvalue of *RT*. Consequently, the DR matrix becomes negative overall, and the state that should be emphasized the most ends up having the most negative value.

This has two drawbacks: (1) it is biologically implausible to observe negative values in the place field, and (2) it causes the agent to avoid states that should be desirable.

One might consider dynamically adjusting the maximum element of *R* to balance the eigenvalues of *RT* just below 1, thereby amplifying subgoals for effective navigation. However, this approach presents significant challenges. Calculating the maximum eigenvalue of *RT* is nontrivial, especially when the transition matrix is unknown and the DR matrix is updated using the Woodbury identity.

Therefore, it is more plausible to bound *r*_*N*_ *<* 0 ∀*r*_*N*_, ensuring that every element of *R* is less than one. This guarantees that the maximum eigenvalue of *RT* remains below one, maintaining stable and biologically plausible place field representations.

### 5 The impact of eigenvalue distributions on structural encoding

The spectral properties of eigenvalues and eigenvectors are fundamental to the ability of the successor representation (SR) and transition matrices to encode spatial structures. A key insight is that eigenvectors with lower spatial frequencies should be weighted more heavily, while those with higher spatial frequencies should be attenuated. This principle aligns with the concept of spectral regularization proposed by Stachenfeld et al.^26^, which emphasizes global structural coherence over local noise.

The expected variance of the eigenvector of transition matrix (*q*_*T*_) is defined as^82^:

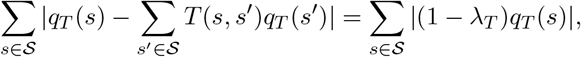

where *λ*_*T*_ denotes the eigenvalue of transition matrix and S denotes a state space. Based on this, we segment the role of eigenvectors with respect to the range of their corresponding eigenvalues. Refer to Supplementary Figure 2 for the visualized examples.

1. *λ*_*T*_ = 1: The eigenvector corresponding to *λ*_*T*_ = 1 represents non-block states where all nodes have identical values, as the expected variance of the eigenvector is 0. This eigenvector lacks spatial differentiation and is therefore not meaningful for structural analysis. It can be neglected when calculating diffusion or successor distances, as it does not distinguish between non-block states.
2. 0 ≤ *λ*_*T*_ *<* 1: Eigenvectors in this range exhibit grid cell-like representations, periodically encoding spatial structures. As the eigenvalue decreases, the expected variance also increases, meaning that the associated spatial frequency increases. Therefore, the eigenvectors with larger eigenvalues tend to capture the global structure, whereas those with smaller eigenvalues tend to capture structural details. The eigenvector with the highest eigenvalue in this range is known as the *Fiedler vector* ^67^, which divides the environmental structure into two regions.
3. *λ* _*T*_ = 0: Eigenvectors with *λ*_*T*_ = 0 are specific to block states. Since _*s*_′_∈S_ *T* (*s, s*^′^)*q*_*T*_ (*s*^′^) = 0 ∀*s* while maintaining ||*q*_*T*_ ||_2_ = 1, these eigenvectors assign a value of 1 to a single block state and 0 to all others, reflecting the inability of these states to propagate transitions.
4. −1 ≤ *λ*_*T*_ *<* 0: In this range, _*s*_′_∈S_ *T* (*s, s*^′^)*q*_*T*_ (*s*^′^) = −|*λ*_*T*_ |*q*_*T*_ (*s*) ∀*s* holds. Therefore, adjacent states have opposite signs, rendering these eigenvectors unsuitable for structural encoding. The alternating sign pattern disrupts meaningful structural representations.

**Figure 1.**
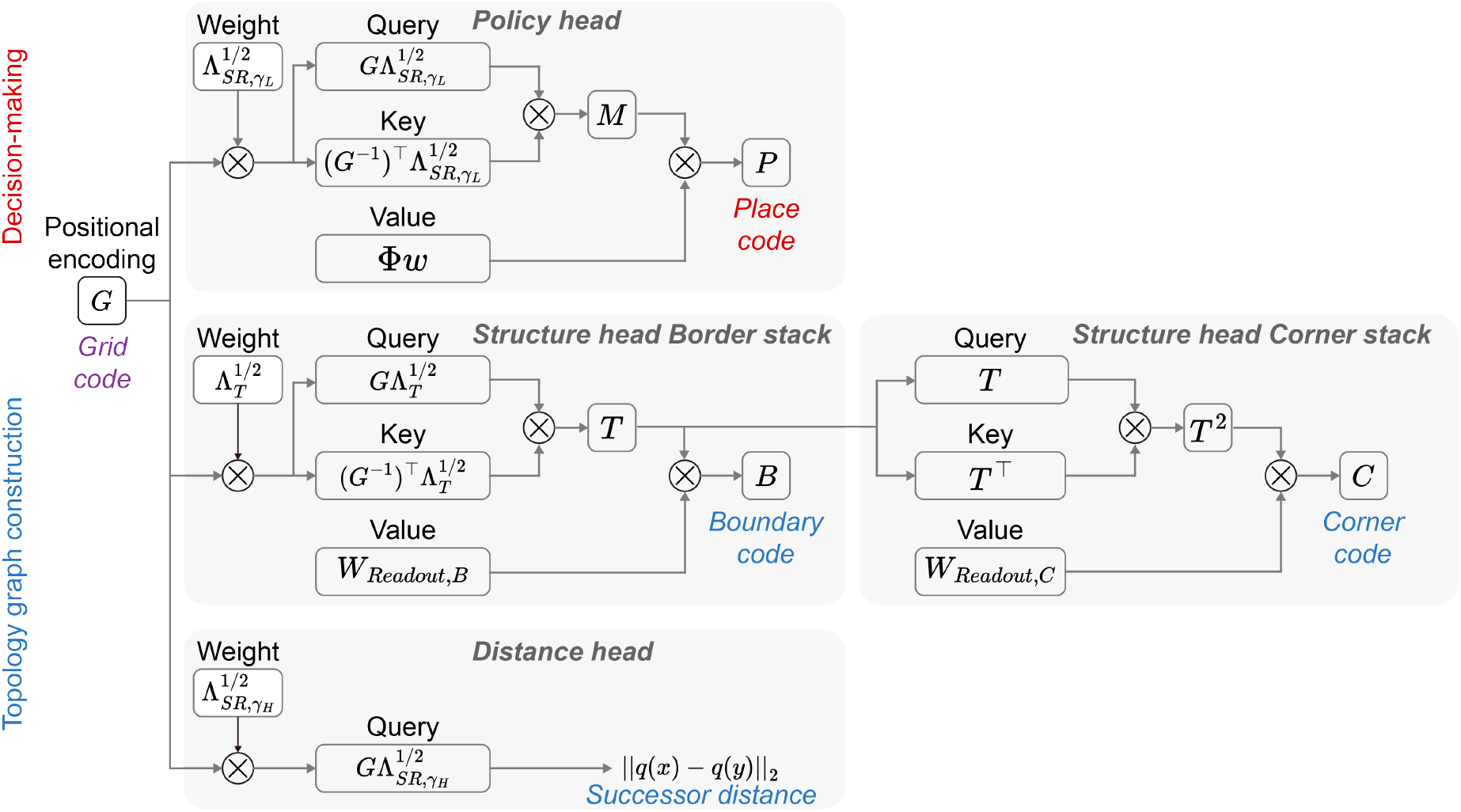
The proposed transformer-like architecture processes a shared grid code with distinct weights across multiple heads to perform specialized roles. The policy head uses self-attention with low-discount SR eigenvalues to calculate place codes guiding immediate navigation targets. The structure head reconstructs the transition matrix to compute a boundary code, which is further refined into a corner code through second-order transitions. The distance head applies self-attention with high-discount SR eigenvalues to measure state dissimilarities, generating edge weights for the topology graph. Together, these outputs construct the topology graph and inform efficient, topology-based decision-making.

**Figure 2.**
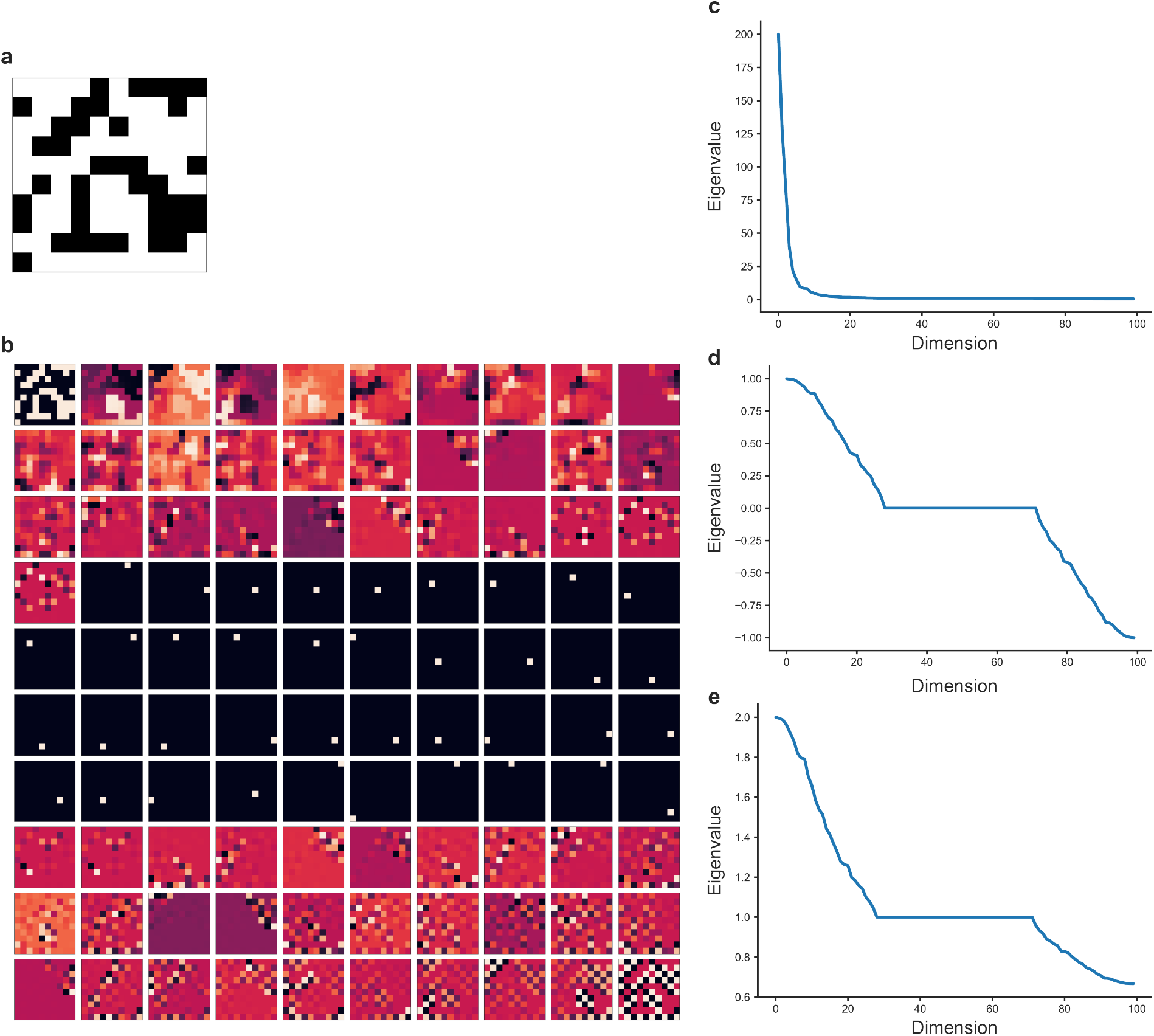
Example eigenvectors and eigenvalues of a maze. **a**, A maze environment. **b**, Eigenvectors of the corresponding transition matrix, with eigenvalues sorted in descending order. **c**, Eigenvalues of the SR with a high discount factor (*γ* = 0.995), sorted in descending order. **d**, Eigenvalues of the transition matrix, sorted in descending order. **e**, Eigenvalues of the SR with a low discount factor (*γ* = 0.1), sorted in descending order.

To understand the distinct characteristics of the eigenvalues in SR and transition matrices, we analyzed three configurations (Supplementary Figure 3b-c):

- (1)SR with a high discount factor (*γ* = 0.995),
- (2)SR with a low discount factor (*γ* = 0.1),
- (3)The transition matrix without discounting.

**Figure 3.**
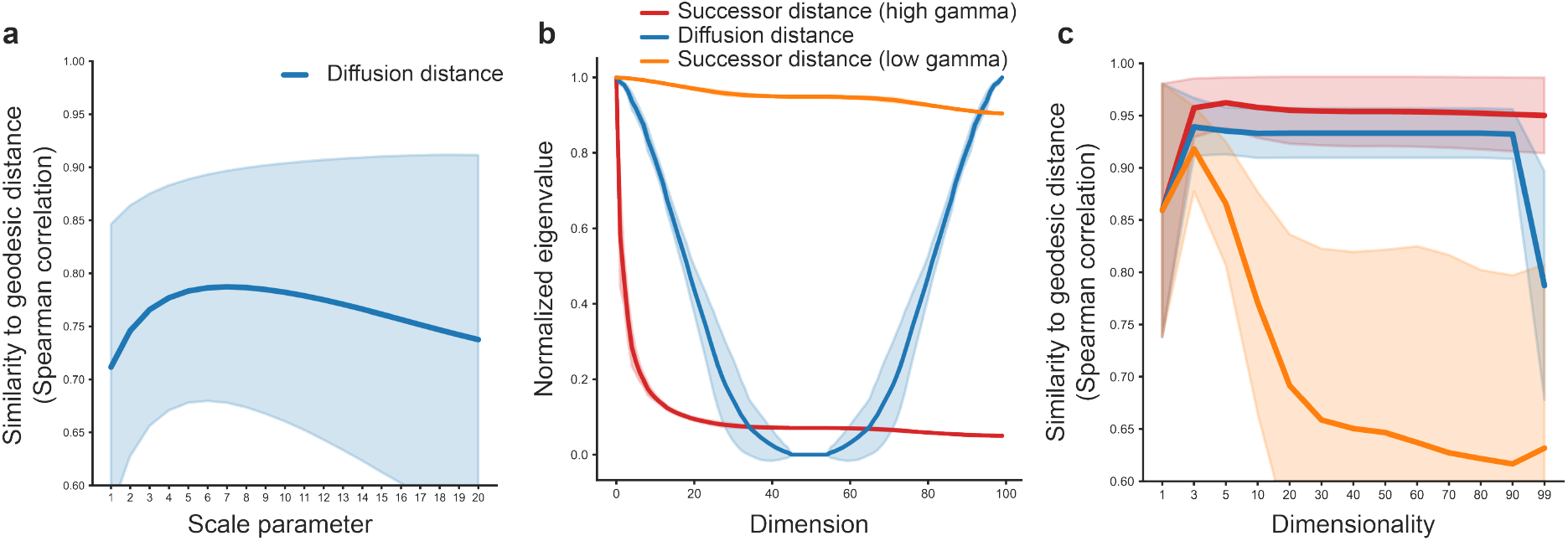
**a**, Structure encoding score (defined as the similarity to geodesic distance) of the diffusion distance as a function of the scale parameter *t*. The best score was achieved at *t* = 7. **b**, Eigenvalues of the SR and transition matrix, corresponding to the values multiplied with the eigenvectors when calculating distances. The eigenvalues of the SR were square-rooted, while those of the transition matrix were taken as absolute values, as the norm used in distance calculations inherently considers absolute magnitudes. To facilitate comparison, each set of eigenvalues was normalized by dividing by its maximum value. **c**, Structure encoding scores for different types of distances. The successor distance with a high discount factor (*γ*) maintains a consistent score as dimensionality increases. In contrast, the diffusion distance does not achieve scores as high as the successor distance with high *γ* when dimensionality is low, due to poor spectral regularization. Furthermore, as dimensionality increases, the score decreases sharply because the eigenvalue distribution amplifies the influence of noisy eigenvectors. Additionally, the successor distance with low *γ* also suffers from reduced structure encoding performance as dimensionality increases, due to poor spectral regularization. Shaded areas represent standard deviations across 1000 randomly generated mazes.

The SR with a high discount factor (*γ* = 0.995) demonstrates superior performance in encoding spatial structures due to effective spectral regularization. By amplifying eigenvectors with larger eigenvalues (corresponding to low spatial frequencies) and suppressing those with smaller eigenvalues (associated with high spatial frequencies), the SR emphasizes global structures while minimizing noise. This mechanism ensures that the resulting representation aligns with the spatial topology of the environment.

In contrast, SR with a low discount factor (*γ* = 0.1) does not sufficiently differentiate between eigenvectors, leading to poor spectral regularization and suboptimal structural encoding. Similarly, the transition matrix exhibits limitations due to its treatment of eigenvalues. In particular, eigenvalues in the negative range produce oscillatory eigenvectors (with high spatial frequencies), which are assigned large weights when squared during the calculation of diffusion distance. These oscillatory components interfere with the representation of spatial structure and introduce noise into the encoding.

Lastly, the transition matrix suppresses eigenvectors with small, nonnegative eigenvalues, although not as effectively as the SR with a high discount factor. As more eigenvectors are involved, the amplification of noisy eigenvectors with high spatial frequencies becomes prominent. Notably, since the norm used in calculating the diffusion distance considers only the absolute magnitude of each axis, contributions from eigenvectors with negative eigenvalues are amplified, thereby reducing the structural encoding score. Although the diffusion distance shows comparable structural encoding performance to the SR with a high discount factor when fewer eigenpairs are involved, its performance deteriorates as more noisy components are included.

### 6 Comparing the generalizability of non-block paths and block objects

In this section, we discuss the generalizability of the predictive object representation (POR)^12^ and justify our focus on the non-block topology of the environment.

First, in real-world scenarios, it is unlikely that the exact same object will reappear. Instead, objects may undergo transformations such as scaling or noise perturbations, similar to the topology-preserving augmentations defined in Methods. Direct object-wise comparison of PORs is challenging because POR matrices vary in size, making it difficult to define meaningful distance metrics between them. Moreover, even if such distances were measurable, there would be no established baseline for interpretation. To address this, we extend the definition of POR to cover the entire environment and compute the L2 norm between PORs from isomorphic and non-isomorphic environment pairs. These isomorphic environment pairs serve as generalized cases of topology-preserving augmentations. As shown in Supplementary Figure 4a, POR distances between isomorphic environments are widely scattered, indicating that POR is less sensitive to topological isomorphisms compared to the TAG grid code.

Second, even when the POR remains unchanged, the overall environmental topology can differ. As illustrated in Supplementary Figure 4b, when two objects are positioned adjacent to one another, they can obstruct topological edges that were previously accessible. This highlights that POR alone does not fully capture the underlying topological structure, reinforcing the importance of considering non-block topology for robust environmental representations.

**Figure 4.**
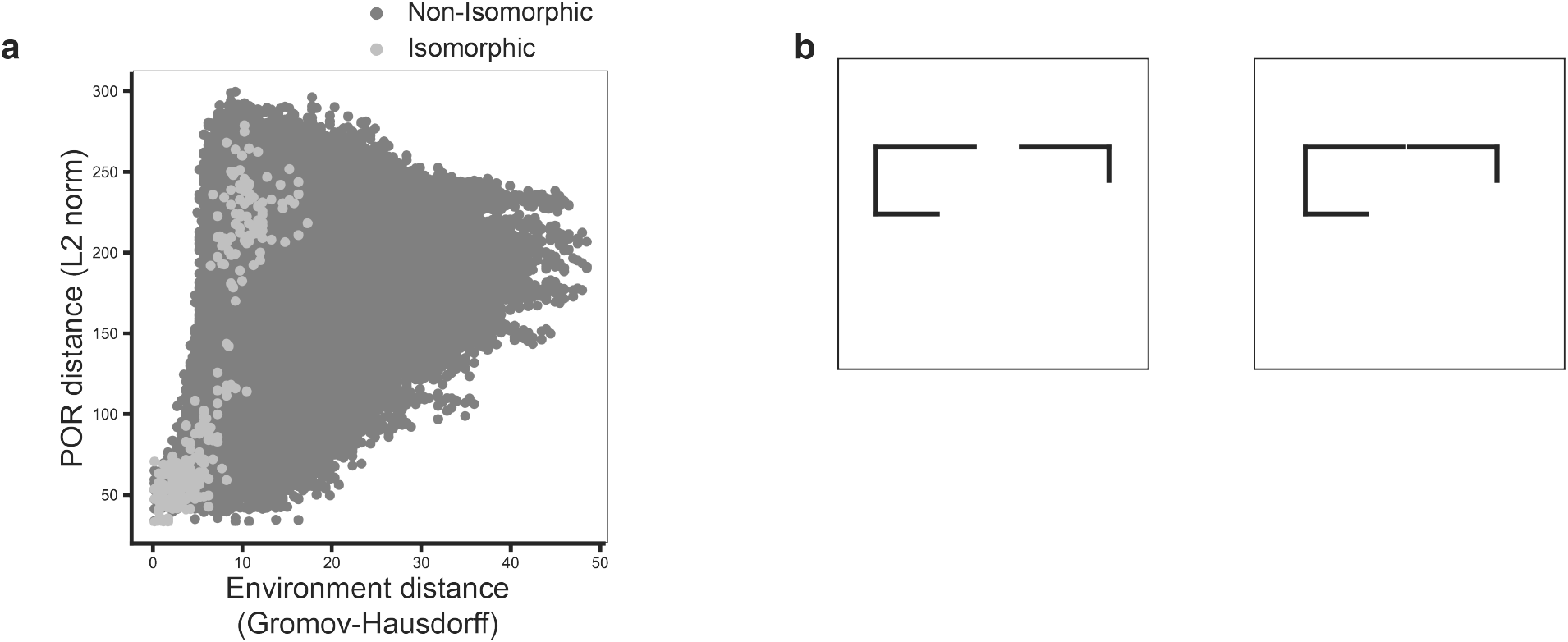
**a**, When the POR is expanded to cover the entire environment, it becomes a metric to capture differences in environmental structures compared to the open field at a state-wise level. The distances between pairwise PORs, each representing a different environment, show that the distances for isomorphic pairs are scattered, implying that state-wise structural comparison is less topology-aware compared to the TAG grid code. To calculate the POR distance, we measured the minimum distance among all the rotation configurations as POR is rotation invariant, only requiring the state reindexing. **b**, Even when the PORs are the same, the topology of the environment may change, resulting in dramatic changes in the overall structure.

**Figure 5.**
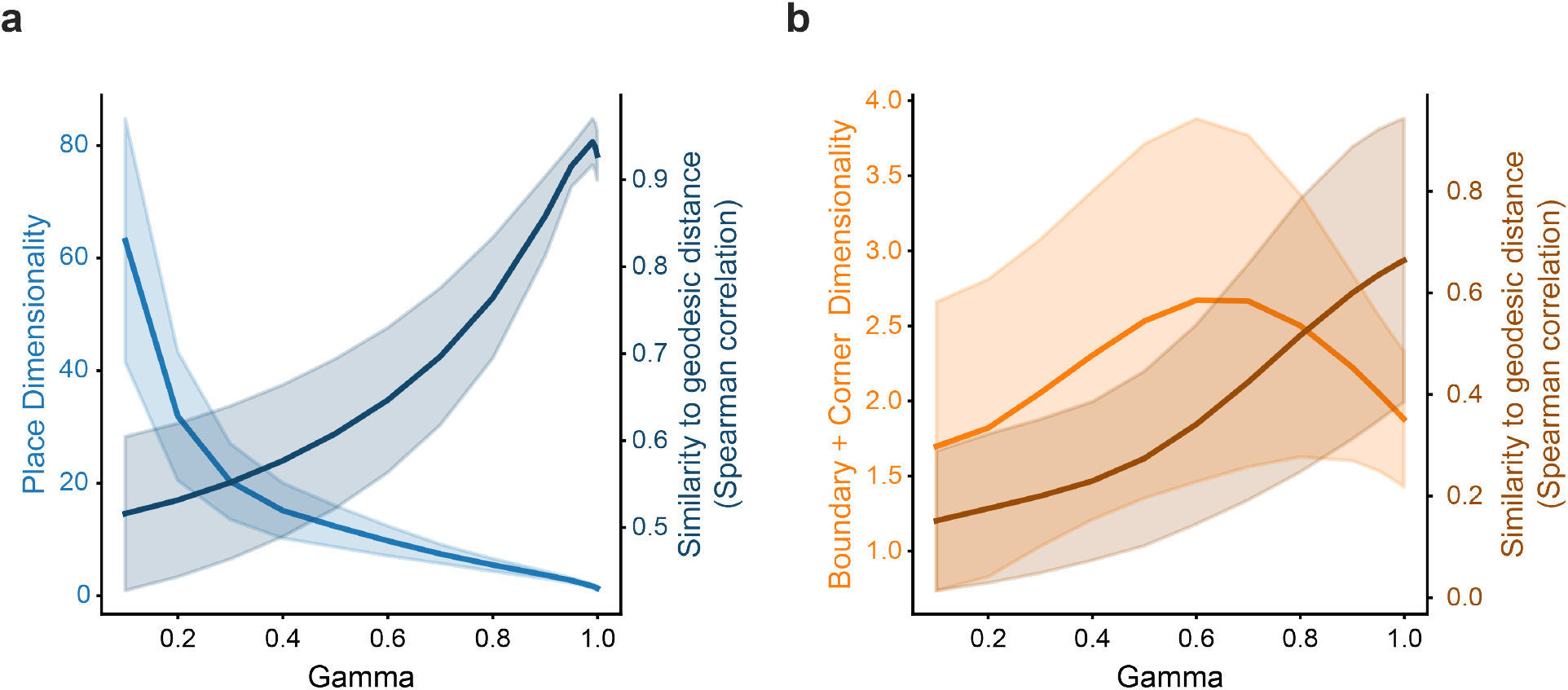
**a**, Dimensionality and similarity to geodesic distance of CA1 neural population in TAG (place code). *γ* = 0.99 yielded the highest similarity to geodesic distance, consistent with the result of Fig. 7c. **b**, Dimensionality and similarity to geodesic distance of subicular neural population in TAG (boundary + corner code). Similar to the result of the CA1, *γ* = 0.99 yielded the highest similarity to geodesic distance.

### 7 Utilizing multiple place fields for subgoal navigation

All SR-related models navigate based on place fields, guiding the agent toward states with higher values. However, relying on a single static place field is insufficient for complex tasks involving multiple subgoals. Instead, multiple place fields are needed to support flexible navigation.

For example, when an agent must first reach a subgoal on a branch and then proceed to the final goal, the place field should initially guide it toward the subgoal and then shift to direct it toward the goal. This requires reorienting the place field’s gradient along the branch to accommodate the new target.

Simply having multiple place fields is not enough; the key is determining their activation order. If multiple place fields are active simultaneously without a defined sequence, the agent may follow the strongest gradient, potentially bypassing intermediate subgoals.

To address this, the agent can either switch between policies tied to specific subgoals, as in SFQL, or maintain multiple place fields with a predefined activation sequence, as in TAG, ensuring the agent follows the intended subgoal order without missing key targets.

### Nonnegativity of SR grid cells

Most studies that frame the eigenvectors of SR as grid cell representations apply a nonnegativity activation function, given that the firing rates of biological cells are inherently nonnegative^11,26,76^. However, in our framework, it is essential to utilize both the positive and negative components of the eigenvectors. This approach is justified for two key reasons: 1. The sign of an eigenvector is not invariant and can always be flipped, making it impossible to fully capture the representation by considering only one side. 2. Topological representations inherently include both positive and negative components, and applying a nonnegativity activation function would result in half of the states being assigned identical distributions, thereby losing critical structural information.

To address this, we propose modeling each eigenvector as two separate grid cells: one representing its positive elements and the other its negative elements. Together, these two grid cells, sharing the same spatial frequency, form a single axis for topological representation.

## Notes

### Competing Interest Statement

The authors have declared no competing interest.

## References

1. Hafting, T., Fyhn, M., Molden, S.Moser, M.-B. & Moser, E. I. Microstructure of a spatial map in the entorhinal cortex. Nature 436, 801–806. issn: 0028-0836 (2005).

2. Okeefe, J. & Dostrovsky, J. The hippocampus as a spatial map. Preliminary evidence from unit activity in the freely-moving rat. Brain Research 34, 171–175. issn: 0006-8993 (1971).

3. Tolman, E. C. Cognitive maps in rats and men. Psychological Review 55, 189–208. issn: 0033-295X (1948).

4. Moser, E. I., Kropff, E. & Moser, M.-B. Place Cells, Grid Cells, and the Brain’s Spatial Representation System. Neuroscience 31, 69–89. issn: 0147-006x (2008).

5. Behrens, T. E. et al. What Is a Cognitive Map? Organizing Knowledge for Flexible Behavior. Neuron 100, 490–509. issn: 0896-6273 (2018).

6. Mattar, M. G. & Daw, N. D. Prioritized memory access explains planning and hippocampal replay. Nature neuroscience 21, 1609–1617. issn: 1097-6256 (2018).

7. Derdikman, D. et al. Fragmentation of grid cell maps in a multicompartment environment. Nature Neuroscience 12, 1325–1332. issn: 1097-6256 (2009).

8. Duvelle, É. et al. Insensitivity of Place Cells to the Value of Spatial Goals in a Two-Choice Flexible Navigation Task. The Journal of Neuroscience 39, 2522–2541. issn: 0270-6474 (2019).

9. Barreto, A. et al. Successor Features for Transfer in Reinforcement Learning. Proceedings of the 31st International Conference on Neural Information Processing Systems, 4058–4068 (2017).

10. Barreto, A., Hou, S., Borsa, D., Silver, D. & Precup, D. Fast reinforcement learning with generalized policy updates. Proceedings of the National Academy of Sciences 117, 30079–30087. issn: 0027-8424 (2020).

11. Piray, P. & Daw, N. D. Linear reinforcement learning in planning, grid fields, and cognitive control. Nature Communications 12, 4942 (2021).

12. Piray, P. & Daw, N. D. Reconciling flexibility and efficiency: medial entorhinal cortex represents a compositional cognitive map. Nature Communications 16, 7444 (2025).

13. Chen, Z., Gomperts, S. N., Yamamoto, J. & Wilson, M. A. Neural Representation of Spatial Topology in the Rodent Hippocampus. Neural Computation 26, 1–39. issn: 0899-7667 (2014).

14. Dabaghian, Y., Mémoli, F., Frank, L. & Carlsson, G. A Topological Paradigm for Hippocampal Spatial Map Formation Using Persistent Homology. PLoS Computational Biology 8, e1002581. issn: 1553-734X (2012).

15. Guo, W., Zhang, J. J., Newman, J. P. & Wilson, M. A. Latent learning drives sleep-dependent plasticity in distinct CA1 subpopulations. Cell Reports 43, 115028. issn: 2211-1247 (2024).

16. Nakai, S., Kitanishi, T. & Mizuseki, K. Distinct manifold encoding of navigational information in the subiculum and hippocampus. Science Advances 10, eadi4471 (2024).

17. Johnson, A. & Redish, A. D. Neural Ensembles in CA3 Transiently Encode Paths Forward of the Animal at a Decision Point. The Journal of Neuroscience 27, 12176–12189. issn: 0270-6474 (2007).

18. Javadi, A.-H. et al. Hippocampal and prefrontal processing of network topology to simulate the future. Nature Communications 8, 14652 (2017).

19. Dabaghian, Y., Brandt, V. L. & Frank, L. M. Reconceiving the hippocampal map as a topological template. eLife 3, e03476 (2014).

20. Barry, C. et al. The Boundary Vector Cell Model of Place Cell Firing and Spatial Memory. Reviews in the Neurosciences 17, 71–98. issn: 0334-1763 (2006).

21. Sun, Y., Nitz, D. A., Xu, X. & Giocomo, L. M. Subicular neurons encode concave and convex geometries. Nature 627, 821–829. issn: 0028-0836 (2024).

22. Park, K., Yeo, Y., Shin, K. & Kwag, J. Egocentric neural representation of geometric vertex in the retrosplenial cortex. Nature Communications 15, 7156 (2024).

23. Cothi, W. & Barry, C. Neurobiological successor features for spatial navigation. Hippocampus 30, 1347–1355. issn: 1050-9631 (2020).

24. Sharp, P. E. Subicular place cells generate the same “map” for different environments: Comparison with hippocampal cells. Behavioural Brain Research 174, 206–214. issn: 0166-4328 (2006).

25. Dayan, P. Improving Generalization for Temporal Difference Learning: The Successor Representation. Neural Computation 5, 613–624. issn: 0899-7667 (1993).

26. Stachenfeld, K. L., Botvinick, M. M. & Gershman, S. J. The hippocampus as a predictive map. Nature Neuroscience 20, 1643–1653. issn: 1097-6256 (2017).

27. McNamee, D. C., Stachenfeld, K. L., Botvinick, M. M. & Gershman, S. J. Flexible modulation of sequence generation in the entorhinal–hippocampal system. Nature Neuroscience 24, 851–862. issn: 1097-6256 (2021).

28. Mahadevan, S. Proto-value functions. Proceedings of the 22nd international Conference on Machine Learning, 553–560 (2005).

29. Mahadevan, S. & Maggioni, M. Proto-Value Functions: A Laplacian Framework for Learning Representation and Control in Markov Decision Processes. J. Mach. Learn. Res. 8, 2169–2231. issn: 1532-4435 (2007).

30. Merris, R. Laplacian graph eigenvectors. Linear Algebra and its Applications 278, 221–236. issn: 0024-3795 (1998).

31. Belkin, M. & Niyogi, P. Laplacian Eigenmaps for Dimensionality Reduction and Data Representation. Neural Computation 15, 1373–1396. issn: 0899-7667 (2003).

32. Amaral, D. G. & Witter, M. P. The three-dimensional organization of the hippocampal formation: a review of anatomical data. Neuroscience 31, 571–591 (1989).

33. Van Strien, N., Cappaert, N. & Witter, M. The anatomy of memory: an interactive overview of the parahippocampal–hippocampal network. Nature reviews neuroscience 10, 272–282 (2009).

34. Canto, C. B., Wouterlood, F. G. & Witter, M. P. What does the anatomical organization of the entorhinal cortex tell us? Neural plasticity 2008, 381243 (2008).

35. Naber, P. & Witter, M. Subicular efferents are organized mostly as parallel projections: a double-labeling, retrograde-tracing study in the rat. Journal of comparative neurology 393, 284–297 (1998).

36. Euler, L. Elementa doctrinae solidorum. Novi commentarii academiae scientiarum Petropolitanae, 109–140 (1758).

37. Russek, E. M., Momennejad, I., Botvinick, M. M., Gershman, S. J. & Daw, N. D. Predictive representations can link model-based reinforcement learning to model-free mechanisms. PLOS Computational Biology 13, e1005768. issn: 1553-734X (2017).

38. Botvinick, M. et al. Reinforcement Learning, Fast and Slow. Trends in Cognitive Sciences 23, 408– 422. issn: 1364-6613 (2019).

39. Wang, J. X. Meta-learning in natural and artificial intelligence. Current Opinion in Behavioral Sciences 38, 90–95. issn: 2352-1546 (2021).

40. Meister, M. Learning, fast and slow. Current Opinion in Neurobiology 75, 102555. issn: 0959-4388 (2022).

41. Butola, T. et al. Hippocampus shapes entorhinal cortical output through a direct feedback circuit. Nature Neuroscience 28, 811–822. issn: 1097-6256 (2025).

42. Gershman, S. J. The Successor Representation: Its Computational Logic and Neural Substrates. Journal of Neuroscience 38, 7193–7200. issn: 0270-6474 (2018).

43. Vaswani, A. et al. Attention Is All You Need. Advances in Neural Information Processing Systems (2017).

44. Barry, C., Hayman, R., Burgess, N. & Jeffery, K. J. Experience-dependent rescaling of entorhinal grids. Nature Neuroscience 10, 682–684. issn: 1097-6256 (2007).

45. Machado, M. C. et al. Eigenoption Discovery through the Deep Successor Representation. International Conference on Learning Representations (2018).

46. Shi, J. & Malik, J. Normalized cuts and image segmentation. IEEE Transactions on Pattern Analysis and Machine Intelligence 22, 888–905. issn: 0162-8828 (2000).

47. Ho, M. K. et al. People construct simplified mental representations to plan. Nature 606, 129–136. issn: 0028-0836 (2022).

48. Whittington, J. C. et al. The Tolman-Eichenbaum Machine: Unifying Space and Relational Memory through Generalization in the Hippocampal Formation. Cell 183, 1249–1263.e23. issn: 0092-8674 (2020).

49. Whittington, J. C. R., Warren, J. & Behrens, T. E. J. Relating transformers to models and neural representations of the hippocampal formation. International Conference on Learning Representations (2022).

50. Kreuzer, D., Beaini, D., Hamilton, W. L., Létourneau, V. & Tossou, P. Rethinking Graph Transformers with Spectral Attention. Advances in Neural Information Processing Systems (2021).

51. Dwivedi, V. P. & Bresson, X. A Generalization of Transformer Networks to Graphs. AAAI’21 Workshop on Deep Learning on Graphs: Methods and Applications (2021).

52. Pan, Y. et al. Exploring Global Diversity and Local Context for Video Summarization. arXiv (2022).

53. McCarter, C. Inverse distance weighting attention. arXiv (2023).

54. Schapiro, A. C., Rogers, T. T., Cordova, N. I., Turk-Browne, N. B. & Botvinick, M. M. Neural representations of events arise from temporal community structure. Nature Neuroscience 16, 486– 492. issn: 1097-6256 (2013).

55. Warren, W. H., Rothman, D. B., Schnapp, B. H. & Ericson, J. D. Wormholes in virtual space: From cognitive maps to cognitive graphs. Cognition 166, 152–163 (2017).

56. Garvert, M. M., Dolan, R. J. & Behrens, T. E. A map of abstract relational knowledge in the human hippocampal–entorhinal cortex. eLife 6, e17086 (2017).

57. Peer, M., Brunec, I. K., Newcombe, N. S. & Epstein, R. A. Structuring Knowledge with Cognitive Maps and Cognitive Graphs. Trends in Cognitive Sciences 25, 37–54. issn: 1364-6613 (2020).

58. Kahn, A. E., Bassett, D. S. & Daw, N. D. Trial-by-trial learning of successor representations in human behavior (2024).

59. Wills, T. J., Lever, C., Cacucci, F., Burgess, N. & O’Keefe, J. Attractor Dynamics in the Hippocampal Representation of the Local Environment. Science 308, 873–876. issn: 0036-8075 (2005).

60. Kinsky, N. R., Sullivan, D. W., Mau, W., Hasselmo, M. E. & Eichenbaum, H. B. Hippocampal Place Fields Maintain a Coherent and Flexible Map across Long Timescales. Current Biology 28, 3578–3588.e6. issn: 0960-9822 (2018).

61. Farebrother, J. et al. Proto-Value Networks: Scaling Representation Learning with Auxiliary Tasks. International Conference on Learning Representations (2023).

62. Boccara, C. N., Nardin, M., Stella, F., O’Neill, J. & Csicsvari, J. The entorhinal cognitive map is attracted to goals. Science 363, 1443–1447. issn: 0036-8075 (2019).

## References

63. Zhang, T. Y. & Suen, C. Y. A fast parallel algorithm for thinning digital patterns. Communications of the ACM 27, 236–239. issn: 0001-0782 (1984).

64. Lin, C. et al. Point2Skeleton: Learning Skeletal Representations from Point Clouds. 2021 IEEE/CVF Conference on Computer Vision and Pattern Recognition (CVPR) 00, 4275–4284 (2021).

65. Coifman, R. R. & Lafon, S. Diffusion maps. Applied and Computational Harmonic Analysis 21, 5–30. issn: 1063-5203 (2006).

66. Corneil, D. S. & Gerstner, W. Attractor Network Dynamics Enable Preplay and Rapid Path Planning in Maze like Environments. Advances in Neural Information Processing Systems (2015).

67. Fiedler, M. A property of eigenvectors of nonnegative symmetric matrices and its application to graph theory. Czechoslovak Mathematical Journal 25, 619–633. issn: 0011-4642 (1975).

68. Mémoli, F. Gromov–Wasserstein Distances and the Metric Approach to Object Matching. Foundations of Computational Mathematics 11, 417–487. issn: 1615-3375 (2011).

69. Gromov, M. Structures métriques pour les variétés riemanniennes. Textes Math. 1 (1981).

70. Raisi, Z., Naiel, M. A., Fieguth, P., Wardell, S. & Zelek, J. 2D Positional Embedding-based Transformer for Scene Text Recognition. Journal of Computational Vision and Imaging Systems 6, 1–4 (2021).

71. Shapley, L. S. Notes on the n-person game—ii: The value of an n-person game (1951).

72. Gower, J. C. & Dijksterhuis, G. B. Procrustes Problems. OUP Oxford, 146–155 (2004).

73. Wang, C. & Mahadevan, S. Manifold alignment using Procrustes analysis. Proceedings of the 25th international conference on Machine learning - ICML ‘08, 1120–1127 (2008).

74. Sanfeliu, A. & Fu, K.-S. A distance measure between attributed relational graphs for pattern recognition. IEEE Transactions on Systems, Man, and Cybernetics SMC-13, 353–362. issn: 0018-9472 (1983).

75. Rubin, A. et al. Revealing neural correlates of behavior without behavioral measurements. Nature communications 10, 4745 (2019).

## References

76. Stachenfeld, K. L., Botvinick, M. & Gershman, S. J. Design Principles of the Hippocampal Cognitive Map. Advances in Neural Information Processing Systems 27 (ed {Weinberger Z. Ghahramani and M. Welling and C. Cortes and N. Lawrence and K.Q.}) (2014).

77. Keck, J., Barry, C., Doeller, C. F. & Jost, J. Impact of symmetry in local learning rules on predictive neural representations and generalization in spatial navigation. PLOS Computational Biology 21, e1013056 (2025).

78. Machado, M. C., Barreto, A. & Precup, D. Temporal Abstraction in Reinforcement Learning with the Successor Representation. arXiv (2021).

79. Ghosh, D. & Bellemare, M. G. Representations for Stable Off-Policy Reinforcement Learning. International Conference on Machine Learning. Proceedings of Machine Learning Research 119, 3556– 3565 (2020).

80. Chandak, Y. et al. Representations and Exploration for Deep Reinforcement Learning using Singular Value Decomposition. International Conference on Machine Learning (2023).

81. Blier, L., Tallec, C. & Ollivier, Y. Learning Successor States and Goal-Dependent Values: A Mathematical Viewpoint. arXiv (2021).

82. Lyle, C., Rowland, M., Dabney, W., Kwiatkowska, M. & Gal, Y. Learning Dynamics and Generalization in Reinforcement Learning. International Conference on Machine Learning (2022).

